# Hierarchical Breakdown of RNA Structure Prediction in CASP16: From Reliable Local Helices to Speculative Multimer Assembly

**DOI:** 10.64898/2026.04.22.720187

**Authors:** Chandran Nithin, Smita P Pilla, Sebastian Kmiecik

## Abstract

CASP16 provided a community-wide benchmark for assessing RNA structure prediction, including the first large-scale blind assessment of RNA–RNA multimer prediction. CASP16 results showed that accurate three-dimensional modeling, especially for RNA– RNA multimers, remains a major challenge across the field. In this work, we use the submissions of our group (LCBio) as a diagnostic case study to examine the current limits of RNA structure prediction. In the official CASP16 best-of-submitted-models analysis, our workflow ranked first in the RNA–RNA multimer category and remained competitive for monomers. This makes the submitted model set useful for examining why high-ranking multimer predictions can still deviate substantially from experimental structures. We combine hierarchical analysis with representative case studies to connect this field-wide limitation to specific structural failure modes, showing that prediction accuracy decreases from relatively reliable canonical base-pairing and local helical organization to less reliable non-canonical interactions, stacking geometry, tertiary motifs, and assembly-level features. In RNA–RNA multimers, errors in monomer structure can combine with uncertainty in interface geometry and model selection, reducing the accuracy of the assembled complexes. These findings point to monomer structure accuracy, interface modeling, and model selection as key areas for improving RNA–RNA multimer prediction.

**Graphical Abstract:** 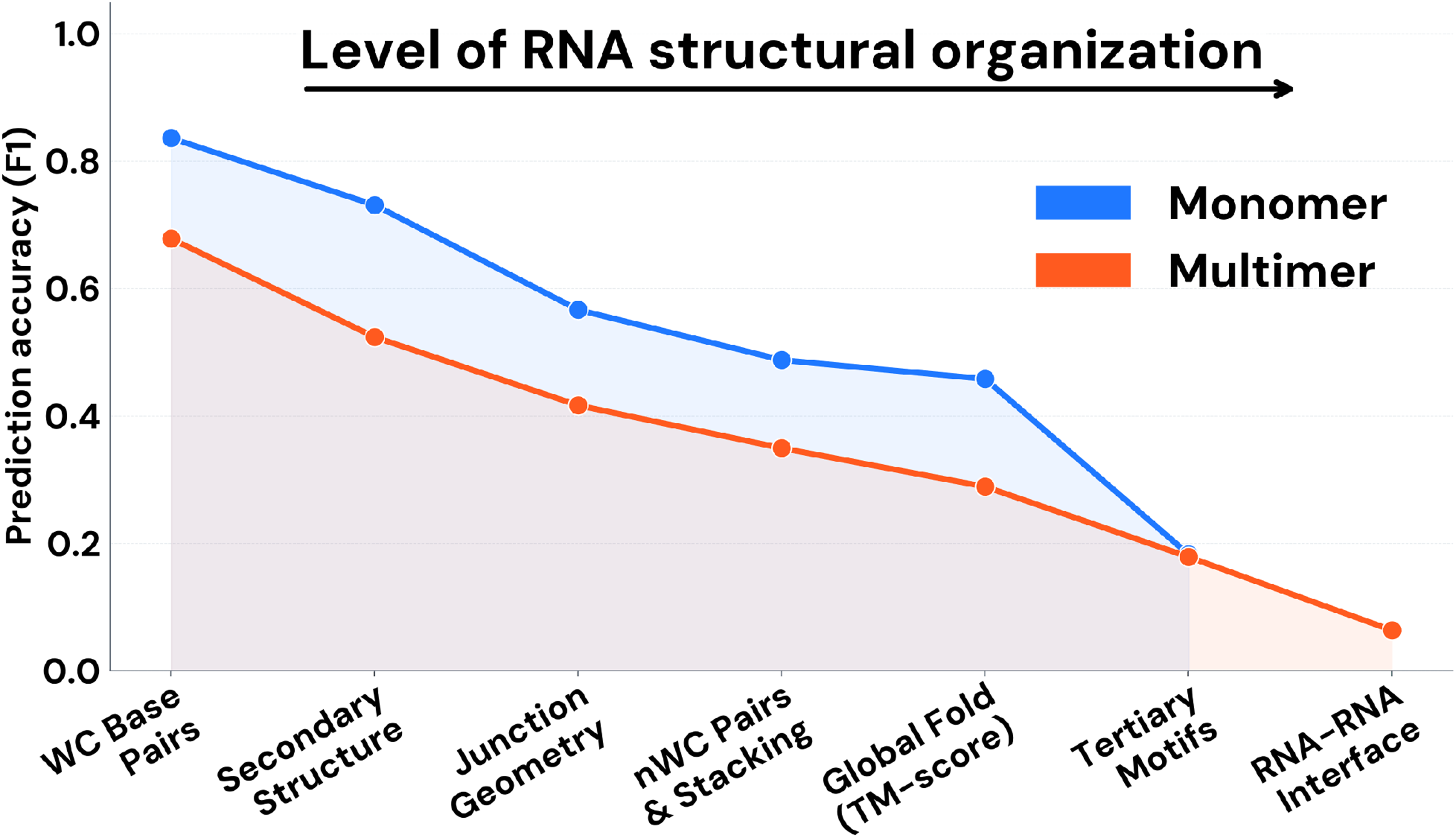

**Key Points:**

1. Using the LCBio CASP16 submissions as a target-level case study, we place known RNA prediction limitations in structural context, especially for RNA–RNA multimers.
2. Canonical helices are recovered more reliably than tertiary motifs, long-range contacts, and RNA–RNA interfaces.
3. Non-canonical interactions and stacking geometry remain difficult to recover accurately in current RNA modeling workflows.
4. The LCBio workflow could generate strong RNA–RNA multimer candidates, but selecting the best model during blind prediction remained difficult.
5. Many CASP16 RNA–RNA multimer models capture only the approximate organization of the complex, not precise atomic structure.

## Introduction

The functional diversity of RNA molecules, extending from their canonical roles in the central dogma to complex regulatory and catalytic activities, is intimately linked to their ability to fold into specific three-dimensional structures (Mortimer, Kidwell, and Doudna 2014, Cao X *et al*. 2024). While protein structure prediction has undergone a profound transformation driven by deep learning (Meng *et al*. 2025, Niazi 2025), de novo and ab initio prediction of RNA architecture remains fundamentally limited (Bahai *et al*. 2024, Mukherjee *et al*. 2024, Nithin *et al*. 2024, Bernard Clément *et al*. 2025, Wang J *et al*. 2025, Elofsson 2026, Ludaic and Elofsson 2026), despite recent methodological advances (Pearce, Omenn, and Zhang 2022, Justyna, Antczak, and Szachniuk 2023, Sato and Hamada 2023, Wang W *et al*. 2023, Abramson *et al*. 2024, Shen *et al*. 2024, Kagaya *et al*. 2025, Zhang S *et al*. 2026). The intrinsic flexibility of the sugar–phosphate backbone, the prevalence of non-canonical base pairing, and the strong dependence on solvent- and ion-mediated electrostatics create a vast and rugged conformational landscape that continues to test the limits of current computational methods (Justyna, Antczak, and Szachniuk 2023). As a result, RNA structure prediction remains uneven across hierarchical levels of organization, with modeling fidelity often declining from local features to global architectures.

The Critical Assessment of Protein Structure Prediction (CASP), a biennial community-wide experiment, serves as a central benchmark for evaluating and driving progress in biomolecular modeling. The recent inclusion and expansion of RNA and RNA-containing complexes as prediction targets within CASP now provide an objective framework for assessing the state of the art in RNA structure prediction (Kryshtafovych *et al*. 2023). Notably, CASP16 represented the first large-scale blind benchmark to explicitly assess RNA–RNA multimer structure prediction (Wu *et al*. 2023), introducing 11 RNA-only multimer targets alongside 35 nucleic acid monomer targets (Kryshtafovych *et al*. 2026).

Official assessment papers summarize overall performance and identify broad methodological trends (Kretsch, Albrecht *et al*. 2026, Kretsch, Hummer *et al*. 2026, Westhof *et al*. 2026). In CASP16, prediction methods fell into a limited number of broad methodological classes (Kagaya *et al*. 2026, Kong *et al*. 2026, Xiao, Shi, and Huang 2026, Zhang S *et al*. 2026)—including deep learning–based approaches, physics-informed sampling, and hybrid workflows—yet the official assessments converge on common structural limitations, particularly in modeling non-canonical interactions, multi-helix junctions, and RNA–RNA interfaces. While emerging quality assessment frameworks are beginning to provide more nuanced measures of RNA model accuracy (Bernard Clement *et al*. 2024), these community-wide evaluations necessarily abstract away the specific modeling decisions and structural failure modes encountered by individual groups.

From a participant’s perspective, the practical limitations of current methods often become visible first in the specific modeling decisions—the “hand of the modeler”— including how secondary structure consensus is constructed, how homology information is interpreted, and how RNA–RNA interfaces are evaluated. This viewpoint makes it possible to connect concrete computational strategies with their structural outcomes and to pinpoint where predictive fidelity degrades from reliable local features to increasingly speculative global assemblies. In particular, this perspective highlights how errors in local and tertiary features can propagate into uncertain global assemblies, especially for RNA– RNA multimers.

In this work, we analyze the current limits of RNA structure prediction using our group’s (LCBio, G189) CASP16 results as a case study. In the official CASP16 best-of-submitted-models analysis, our workflow achieved the top ranking in the RNA–RNA multimer category and remained competitive for monomeric RNA targets. In this analysis, the top-ranking result was based on the best model selected from our submitted set after the experimental structures were available. It therefore reflects the ability of the workflow to generate good candidate models, but not necessarily to identify the best one during the blind prediction stage. Consistent with this distinction, LCBio ranked lower under CASP16 rankings based on submitted model order or average model quality. By examining where this model set succeeds or fails, we show that current methods are relatively reliable for local secondary structure features but become increasingly uncertain for more complex tertiary and assembly-level features. We therefore suggest that contemporary multimer models should often be interpreted as coarse-grained organizational hypotheses rather than precise atomic structures.

## 2. Methods

### 2.1 A Consensus-Based Pipeline for RNA Secondary Structure Prediction

RNA secondary structure prediction was performed using a consensus-based framework designed to integrate complementary sources of structural information while explicitly preserving structural uncertainty. As summarized in Figure 1 (Step 1), the pipeline combined de novo secondary structure prediction, pseudoknot-aware modeling, homology-informed refinement, and manual curation to generate secondary-structure restraints for downstream three-dimensional modeling.

**Figure 1.**
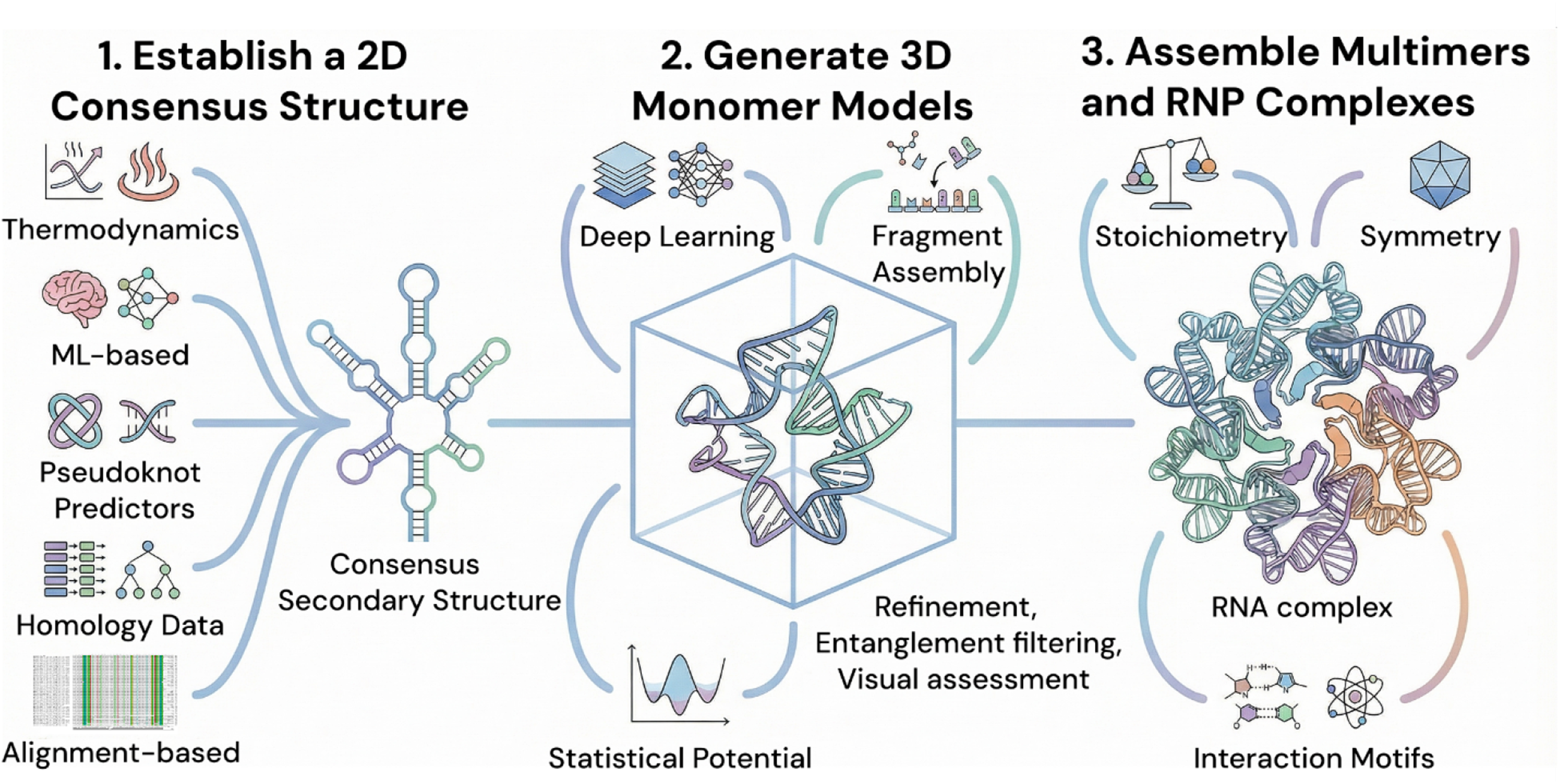
Three-step overview of the LCBio (189) modeling workflow used in CASP16. In Step 1, RNA secondary structure restraints are derived from a consensus of complementary prediction methods and are refined using homologous information from Rfam, RNAcentral, and the Protein Data Bank, with manual curation used to preserve features such as pseudoknots (Supplementary Methods S1). In Step 2, candidate RNA 3D monomer models are generated using multiple complementary approaches and filtered and selected on the basis of structural plausibility, agreement across models, refinement, and lack of entanglements (Supplementary Methods S2.1–S2.2). In Step 3, higher-order assemblies are constructed as required, with RNA–RNA multimer modeling guided by stoichiometry, symmetry, and interface considerations, and RNA–protein complex assembly guided by structural templates and conserved interaction motifs (Supplementary Methods S2.3–S2.4). Alt text: Schematic of a three-step RNA structure modeling workflow. Consensus secondary structures are established from complementary prediction methods and homology information, 3D monomer models are generated and refined, and higher-order RNA multimers and RNA–protein complexes are assembled using stoichiometry, symmetry, structural templates, and interaction motifs.

For each target, an ensemble of candidate secondary structures was generated using multiple methods spanning thermodynamic, probabilistic, and pseudoknot-aware approaches (Do, Woods, and Batzoglou 2006, Lu ZJ, Gloor, and Mathews 2009, Sato *et al*. 2009, 2011, Bellaousov and Mathews 2010, Lorenz *et al*. 2011, Ali, Mittal, and Mathews 2023); the complete list of tools and parameters is given in Supplementary Methods S1.1. When homologous sequences or structurally related RNAs were identified, this information was used to refine the ensemble and incorporate conserved base-pairing patterns supported by external annotations or known structures (Ontiveros-Palacios *et al*. 2025, Consortium *et al*. 2026, Vallat *et al*. 2026). Additional local-stability analyses (ScanFold 2.0 and RNALfold) were incorporated for longer RNAs where sequence length made them informative; parameters are given in Supplementary Methods S1.3 (Hofacker, Priwitzer, and Stadler 2004, Washietl and Hofacker 2004, Bernhart *et al*. 2008, Gruber *et al*. 2009, Nawrocki and Eddy 2013, Andrews, Roche, and Moss 2018, Andrews *et al*. 2022, Zhang C, Zhang, and Pyle 2023).

All candidate structures were converted to dot-bracket notation and aggregated into a unified ensemble. Consensus base-pairing restraints were then derived from base-pair occurrence frequencies across this ensemble. To encode uncertainty explicitly, we retained two representative restraint sets for downstream three-dimensional modeling: a permissive set capturing moderately supported interactions and a stringent set retaining only universally supported base pairs. In cases of mutually exclusive pairing patterns, alternative restraint sets were preserved as separate structural hypotheses.

Following consensus construction, candidate structures were manually inspected and curated using two-dimensional representations. Manual intervention was restricted to four defined stages: adjustment of template-informed regions, alignment cleaning, inspection of consensus structures, and targeted correction of implausible base-pairing patterns. The finalized consensus structures were converted into base-pairing restraints and used as primary inputs for subsequent three-dimensional modeling. Detailed tool lists, thresholds, and target-specific decision rules are provided in Supplementary Methods S1.

### 2.2 A Consensus-Based Pipeline for RNA Tertiary and Complex Prediction

Consensus-derived secondary structure restraints were used as primary inputs for three-dimensional (3D) structure prediction. As outlined in Figure 1 (Steps 2 and 3), candidate monomer models were generated using multiple complementary approaches and then used as starting points for RNA–RNA multimer and RNA–protein assembly where required, through a combination of refinement, filtering, and expert-guided model selection.

To explore conformational space broadly and reduce dependence on any single modeling strategy, we generated candidate monomer models using multiple complementary approaches spanning deep learning–based predictors, fragment-assembly methods, and simulation-based strategies (Cao S and Chen 2005, 2011, Popenda *et al*. 2012, Boniecki *et al*. 2016, Xu and Chen 2018, Manfredonia *et al*. 2020, Watkins Andrew Martin, Rangan, and Das 2020, Wirecki *et al*. 2020, Xiong *et al*. 2021, Li *et al*. 2022, Pearce, Omenn, and Zhang 2022, Sarzynska *et al*. 2023, Wang W *et al*. 2023, Watkins Andrew M. and Das 2023, Abramson *et al*. 2024, Shen *et al*. 2024, Kagaya *et al*. 2025); the list of tools, parameters, and model counts is given in Supplementary Methods S2.1. For methods that allowed it, modeling was performed both with and without secondary structure restraints, and models produced under different conditions were treated as independent structural hypotheses.

Across all methods, multiple candidate models were generated for each target and filtered using a combination of automated and manual criteria. Refinement was applied uniformly prior to final evaluation (Stasiewicz *et al*. 2019, Luwanski *et al*. 2022), and models with non-physical geometry (defined in Supplementary Methods S2.2) were removed. Final submitted models were selected to maximize both structural plausibility and conformational diversity across the candidate pool.

For RNA–RNA multimer targets, higher-order assemblies were generated by evaluating alternative stoichiometries and arranging monomer models into candidate complexes using structural plausibility, symmetry considerations, and available template information where relevant. For RNA–protein complexes, conserved interaction motifs and homologous structural information were used to guide interface placement and assembly (Barik *et al*. 2015, 2016, Nithin, Mukherjee, and Bahadur 2019, Mukherjee and Nithin 2022). In both cases, expert-guided inspection played an important role in filtering and selecting the final submitted models. The four defined stages at which manual intervention was applied during 3D refinement, filtering, and assembly are explained in Supplementary Methods S2.

This workflow reflects the practical modeling strategy used in CASP16: broad exploration at the monomer level, followed by increasingly selective assembly and model curation at higher levels of organization. Detailed modeling protocols, tool-specific settings, model counts, simulation parameters, and assembly procedures are provided in Supplementary Methods S2.

### 2.3 Official CASP16 metrics and post-CASP structural analyses

All global and interface evaluation metrics used in this study were taken directly from the official CASP16 assessment (https://predictioncenter.org/download_area/CASP16/) and were not recomputed. Composite performance measures (Z*_monomer_*and Z*_multimer_*) were used to summarize overall model quality across targets. In addition to these benchmark measures, we performed feature-level analyses on the models submitted by the LCBio group in CASP16 to quantify prediction accuracy across successive levels of RNA structural organization. Structural features were extracted from both reference and predicted RNA structures using DSSR (Lu XJ, Bussemaker, and Olson 2015) to ensure consistent annotation across all models. Full definitions of structural features, scoring procedures, and aggregation rules are provided in the Supplementary Methods S3 and Supplementary Table S1.

## 3 Results

### 3.1 CASP16 performance landscape in the context of global benchmarks

The CASP16 experiment provides a comprehensive snapshot of the current state of RNA structure prediction across monomeric RNAs, RNA–RNA multimeric assemblies, and RNA–protein (RNP) complexes, evaluated using a consistent set of global accuracy metrics. As summarized in the official CASP16 assessment (Kretsch, Albrecht *et al*. 2026, Kretsch, Hummer *et al*. 2026, Westhof *et al*. 2026), RNA monomer prediction spans a broad range of quality—from near-random models to structures with recognizable overall topology—whereas RNA–RNA multimer prediction remains largely confined to a low-accuracy regime. In contrast, RNP complexes often show a split between high-accuracy interface recovery, facilitated by structural templates, and lower-accuracy global folding.

To establish a clear reference point for the analyses that follow, Figure 2 presents the distribution of best-achieved TM-scores for each target in CASP16. This view highlights a qualitative separation between monomers and multimers that is implicit in the CASP assessment: while a substantial fraction of monomeric RNAs reach intermediate global similarity to experimental structures (exceeding the TM-score = 0.45 threshold for correct topology), RNA–RNA multimers rarely do so, even for the best-performing models. RNP complexes further demonstrate how the availability of interfacial templates can significantly boost global fold accuracy beyond that of unconstrained multimers.

**Figure 2.**
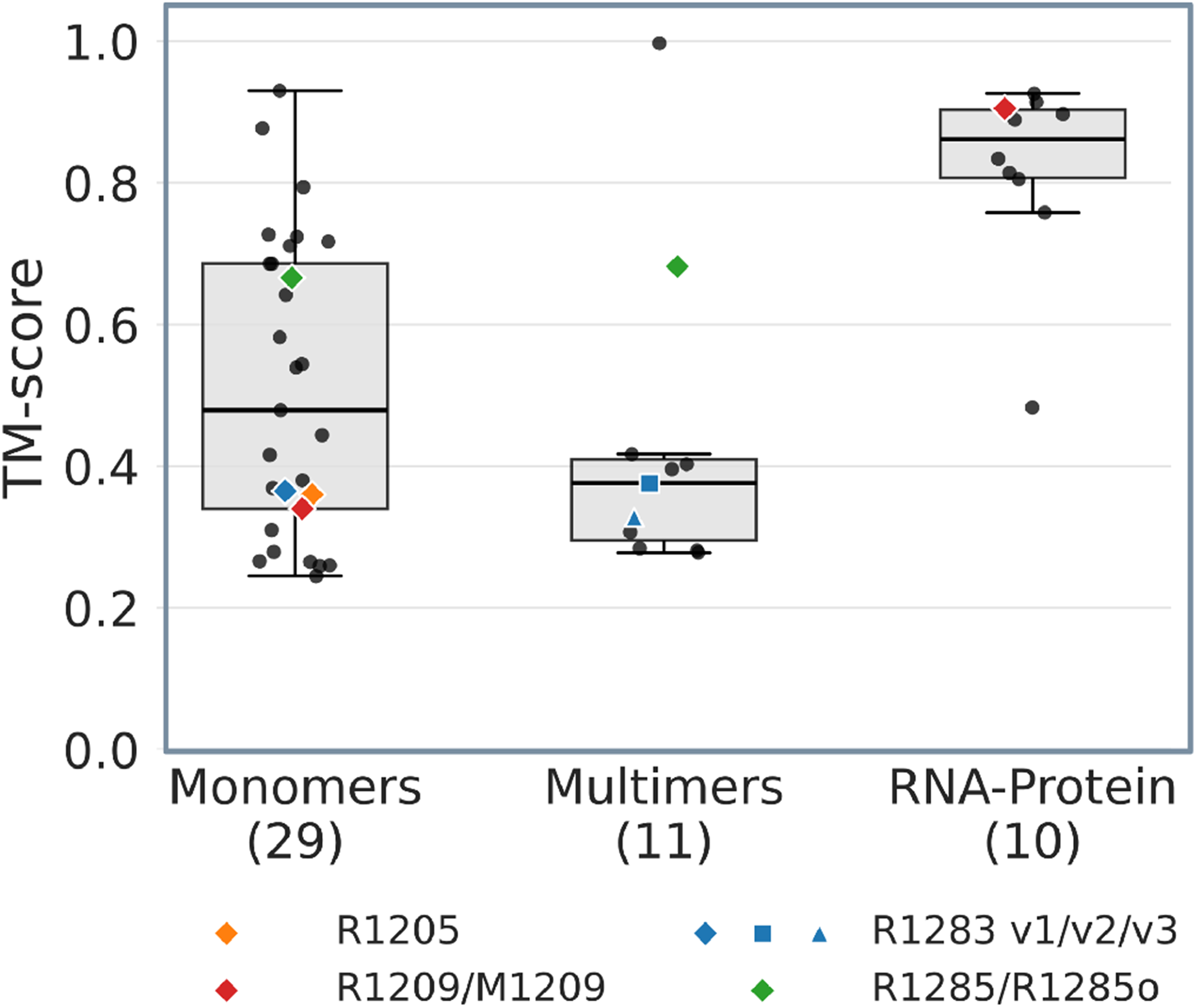
Overall performance of best models across all groups in CASP16 target categories. Boxplots show the distribution of TM-scores for best models across all groups in the Monomer (n = 29), RNA–RNA Multimer (n = 11), and RNA–protein complex (n = 10) categories. Individual targets are shown as grey dots. The dashed horizontal line at TM-score = 0.45 indicates a commonly used threshold for correct global topology. Case studies discussed in the text are highlighted with colored markers: R1205 (orange diamond), R1209/M1209 (red diamond), R1283 variants (blue markers), and R1285/R1285o (green diamonds). The dual placement of the red diamonds illustrates the contrast between the incorrect RNA monomer model (R1209; TM-score < 0.45) and the RNA–protein complex model (M1209; TM-score > 0.8). Alt text: Boxplots compare TM-scores for best models in three CASP16 target categories: monomers, multimers, and RNA–protein complexes. Individual targets are shown as points, with selected case-study targets highlighted by colored symbols. The RNA–protein complex group has the highest overall TM-scores, while monomers show the widest distribution.

This qualitative separation is not specific to CASP16 but is instead largely recapitulated across independent benchmarks. While RNA-Puzzles (Bu *et al*. 2024) and Kaggle (Lee *et al*. 2025) focus primarily on single-chain prediction, CASP16 now extends this picture to RNA–RNA multimers. The general difficulty of modeling non-helical regions, non-canonical contacts, and expert-dependent model selection has been recognized previously. However, RNA–RNA multimer prediction adds a distinct layer of uncertainty, because monomer-level errors interact with assembly assumptions, stoichiometry, interface placement, and submitted-model selection. The value of the present analysis is to place this multimer-specific problem in a target-level structural context using the LCBio CASP16 submissions.

Across these benchmarks, secondary structure and local helical geometry are frequently recovered with reasonable accuracy. In RNA-Puzzles, correct Watson–Crick pairing and stacking interactions are often predicted even when global folds are incorrect, highlighting the contrast between canonical and non-canonical interaction fidelity (Kretsch, Hummer *et al*. 2026). Similarly, as observed for CASP16 monomer targets in this work, a substantial fraction of RNAs reach an intermediate global-fold regime, while local helical elements remain comparatively reliable. These observations are consistent with earlier RNA-Puzzles and CASP conclusions that non-helical nucleotide arrangements, non-canonical contacts, and expert-dependent model selection remain difficult. Here, we use the LCBio CASP16 submissions to connect these established challenges to concrete modeling decisions, feature-level outcomes, and RNA–RNA multimer assemblies.

Importantly, refinement does not resolve these limitations. Systematic benchmarking of molecular dynamics-based refinement in CASP-scale RNA datasets (Nithin, Pilla, and Kmiecik 2025) shows that MD simulations provide, at best, modest improvements for already accurate models and do not reliably correct incorrect tertiary organization or global topology. Poorly predicted models rarely benefit and often deteriorate upon refinement. These observations strengthen the interpretation that current limitations arise not simply from insufficient sampling, but from insufficient information about the higher-order structural features that determine global RNA architecture.

Within this competitive landscape, our group, LCBio (G189), demonstrated robust performance among the 64 participating groups. For RNA monomers, our submissions ranked 14th overall under the official CASP16 best-of-submitted-models analysis (Supplementary Figure S1A). This standing is supported by a target-by-target analysis of relative performance, which shows that our models achieved generally above-average Z-scores across many monomeric targets (Supplementary Figure S2A). Structurally, this places our monomer predictions within the intermediate-quality regime, where correct secondary structure and overall fold topology were successfully recovered for several targets.

In the RNA–RNA multimer category, our pipeline achieved its most notable result, ranking first overall among all participating groups according to the official CASP16 best-of-submitted-models analysis (Supplementary Figure S1B). As noted above, this result reflects the presence of high-scoring models within the submitted set and should be distinguished from prospective model selection during the blind prediction stage. This result identifies our modeling workflow as having strong sampling potential for RNA complexes within the CASP16 experiment. As shown in the weighted heatmap of relative performance (Supplementary Figure S2B), our models often outperformed the community average across both global structural metrics and interface-specific scores (e.g., ICS, IPS, and iLDDT) for many multimer targets. However, this relative success was achieved despite the low absolute accuracy of predicted assemblies across the field, reflecting a community-wide limitation in RNA complex modeling rather than unusually precise atomic predictions. This contrast between best-of-submitted-models performance and limited structural realism motivates the hierarchical analysis that follows.

### 3.2 Hierarchical trends in predictive accuracy

#### 3.2.1 Diagnostic lens and methodological framework

To identify the structural origins of the performance gap described in the previous section, we carried out a high-resolution analysis of the models submitted by the LCBio group (G189). Because this pipeline ranked first in the official CASP16 best-of-submitted-models analysis for RNA–RNA multimers (Supplementary Figure S1B) while ranking 14th in the corresponding monomer analysis (Supplementary Figure S1A), it offers a useful participant-level perspective on the current strengths and limitations of RNA structure prediction. As noted above, this ranking reflects retrospective selection of the strongest model within the submitted set and should be distinguished from prospective model selection during the blind prediction stage. Using these models as a diagnostic lens, we aimed to distinguish broad methodological success from specific structural failure modes across the organizational hierarchy (Figure 3).

**Figure 3.**
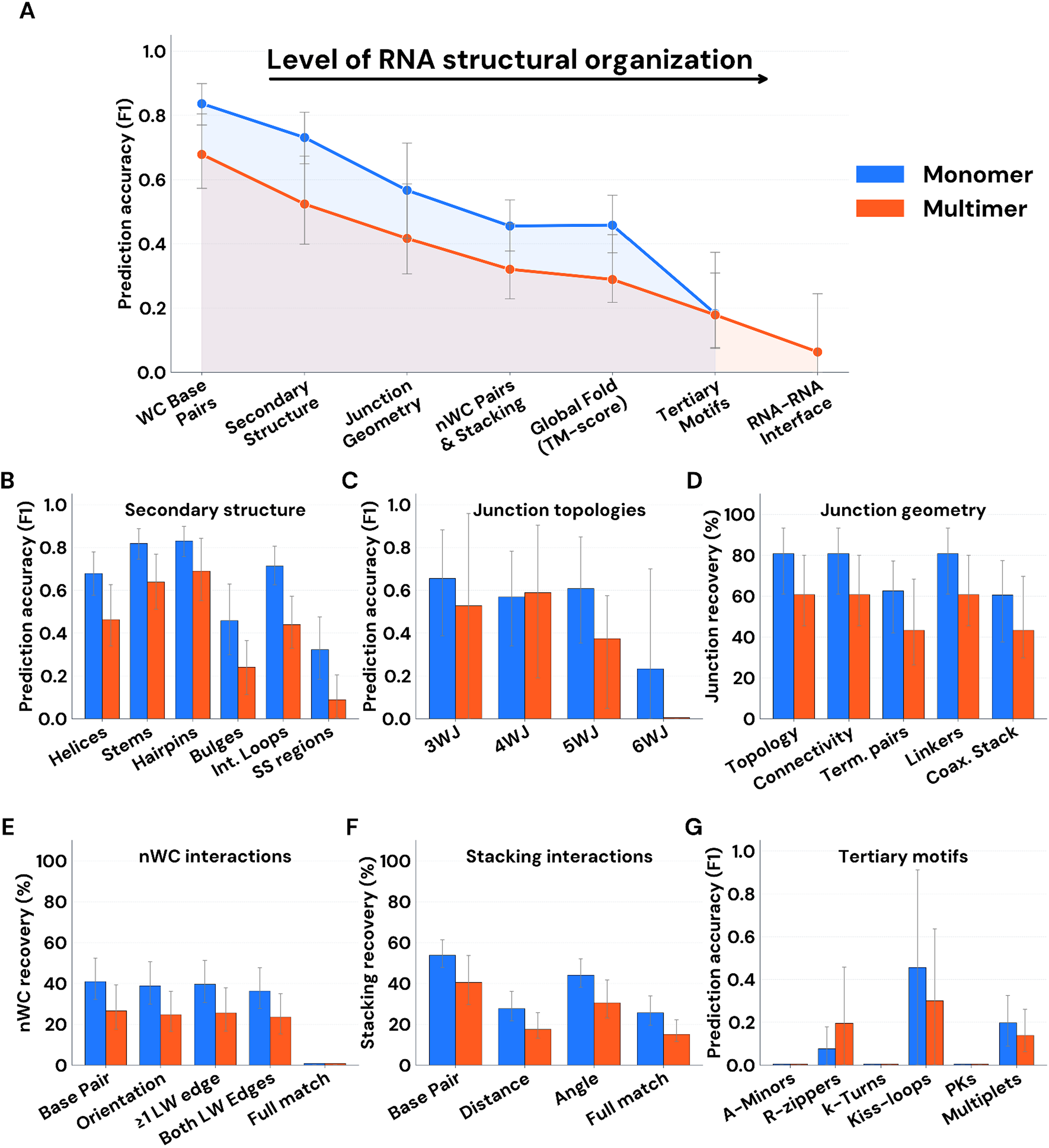
Hierarchical decline in RNA structure prediction accuracy. (A) Global trend of prediction accuracy (F1-score) across hierarchical levels of RNA organization, from local base pairing to global folds and multimer interfaces. (B) Accuracy for specific secondary structure elements, evaluated at the level of base pairs (helices/stems) or residues (loops/segments). (C) Junction topology performance across increasing branching complexity (3WJ to 6WJ) and (D) recovery of junction geometry components, including connectivity and coaxial stacking. (E) Decomposition of non-Watson–Crick (nWC) base pair features and (F) stacking interaction geometry (distance and angular deviation) relative to reference structures. (G) Prediction accuracy for representative tertiary motifs. Results for monomeric and multimeric targets are shown in blue and orange, respectively. Detailed definitions of structural features and scoring procedures are provided in Supplementary Methods S3.2, with relative group performance heatmaps available in Supplementary Figures S1–S2. Error bars in all panels represent 95% bootstrap confidence intervals accounting for non-independence among related multimer targets (see Supplementary Methods S3.3). Alt text: Seven-panel figure showing decreasing RNA structure prediction accuracy as structural organization increases. Panel A shows F1-score trends from base pairing through RNA–RNA interfaces. Panels B–G compare monomer and multimer performance for secondary-structure elements, junction topologies and geometry, non-Watson–Crick interactions, stacking interactions, and tertiary motifs.

The accuracy of structural features was quantified using a standardized annotation pipeline based on DSSR (Lu XJ, Bussemaker, and Olson 2015), with detailed procedures provided in Supplementary Methods S3.2. For each target, a single representative model was selected based on the highest official scores and aligned against the corresponding experimental reference structure. Accuracy for interaction-based features was measured using the F1-score, while global fold accuracy was incorporated via the TM-score. Crucially, our evaluation employed a multi-component scoring scheme: Watson–Crick pairs were evaluated using strict matching, non-Watson–Crick pairs were scored on base identity, cis/trans orientation, and the two Leontis–Westhof edges, and stacking interactions on base identity, inter-planar distance, and angular deviation (Supplementary Methods S3.2).

#### 3.2.2 The hierarchical trend of predictive accuracy

Our analysis indicates that prediction accuracy is not distributed uniformly across structural features. Instead, Figure 3A shows a broad hierarchical trend in which features closely linked to canonical secondary structure are recovered more reliably than features that depend on tertiary organization or higher-order assembly. Monomeric targets generally show higher recovery than RNA–RNA multimers, but the feature-level differences should be interpreted cautiously, especially for multimers, because the CASP16 multimer set is small and includes related targets. We therefore treat these results as descriptive trends within the LCBio model set rather than as evidence for a sharply defined transition at a specific structural level.

#### 3.2.3 Accuracy across secondary structure and local elements

At the baseline of this hierarchy, canonical elements such as helices and stems are recovered with high fidelity (Figure 3B). However, even at this local level, a performance gap emerges in non-canonical regions. Accuracy for internal loops, bulges, and single-stranded (SS) regions is lower, particularly in multimeric targets. This is consistent with our base interaction analysis (Supplementary Figure S3), which shows that while Watson–Crick pairing is often “captured,” the reliability of non-canonical interactions is generally lower across the dataset.

#### 3.2.4 Junction features and three-dimensional organization

Overall, the feature-level analysis supports a broad hierarchical trend in which accuracy is highest for features closely linked to canonical secondary structure and generally decreases for features that depend on more complex tertiary organization or higher-order assembly. Within this trend, Figure 3C shows lower recovery for more complex junction classes, although the small number of highly branched examples limits the strength of this comparison. Figure 3D further indicates that junction-related features are recovered unevenly: connectivity and linker residues show relatively higher recovery, whereas features depending on precise three-dimensional organization, including coaxial stacking, are less consistently recovered. These results indicate increasing uncertainty as prediction moves from local structural features toward higher-order organization and should be interpreted together with the target-level case studies.

#### 3.2.5 Limitations in atomic resolution and tertiary motif recovery

The decline in recovery is also evident at the level of fine-grained interactions. Decomposition of nWC base pairs (Figure 3E) and base stacking geometry (Figure 3F) shows that while base identity is often predicted, the simultaneous recovery of orientation and interacting edges (Full Match) are much less frequent. For nWC interactions, incorrect Leontis–Westhof edges are consistent with the loss of the dense interaction networks required to stabilize complex folds. Similarly, stacking interactions suffer from significant deviations in distance and angle, further destabilizing the predicted architectures. Finally, specialized tertiary motifs (Figure 3G), such as pseudoknots (PKs) and kissing loops, show low recovery in this dataset, with near-zero accuracy for several complex motif classes.

#### 3.2.6 Error propagation and the disconnect between ranking and structural accuracy

Our analysis suggests that multimer prediction can inherit and amplify errors from monomer modeling. This is illustrated by the discrepancy in relative performance: our submitted model set achieved the top ranking in the official CASP16 best-of-submitted-models analysis for RNA–RNA multimers, even though the underlying monomeric subunits clustered with the main cohort of competitive groups rather than the absolute top. This disconnect between ranking and structural accuracy suggests that, at the current state of the art, multimer success depends not only on subunit quality but also on expert-guided assembly and curation. Because several monomeric features, including non-canonical interactions, junction-related features, and tertiary motifs, are recovered less consistently (Figure 3), multimer assembly remains uncertain in many cases. Consequently, many RNA–RNA multimer models in this benchmark are better interpreted as coarse-grained organizational hypotheses than as atomically accurate structures.

### 3.3 Structural drivers of failure: representative case studies

To bridge the gap between our statistical hierarchy and concrete structural outcomes, we analyzed a set of representative targets that exemplify the “capture vs. miss” regime in current RNA modeling. These cases illustrate how the hierarchical breakdown identified in Figure 3 manifests in both monomeric and multimeric systems.

#### 3.3.1 Architectural barriers and tertiary motifs in monomeric RNAs

The failures observed in monomeric targets R1205 and R1209 illustrate the critical transition point where modeling accuracy shifts from reliable 2D components to speculative 3D orientations.

Target R1205 (an exoribonuclease-resistant RNA; xrRNA; PDB ID: 9CFN (Gezelle *et al*. 2025)) highlights the current resolution limit of automated tertiary motif recovery. While our consensus pipeline correctly identified the 2D topology, enforcing the pseudoknot constraint during 3D modeling did not yield native-like structures (Figure 4A). Native xrRNA stabilization depends on a dense network of non-canonical base triples and specialized hairpin conformations. Consistent with the low statistical accuracy for specialized motifs across our entire dataset (Figure 3G), our models for R1205 satisfied the topological constraints at the expense of physically realistic junction geometry. This confirms that correct topology is insufficient for global structural realism when the underlying “tertiary glue”—the non-canonical interactions—is missed.

**Figure 4.**
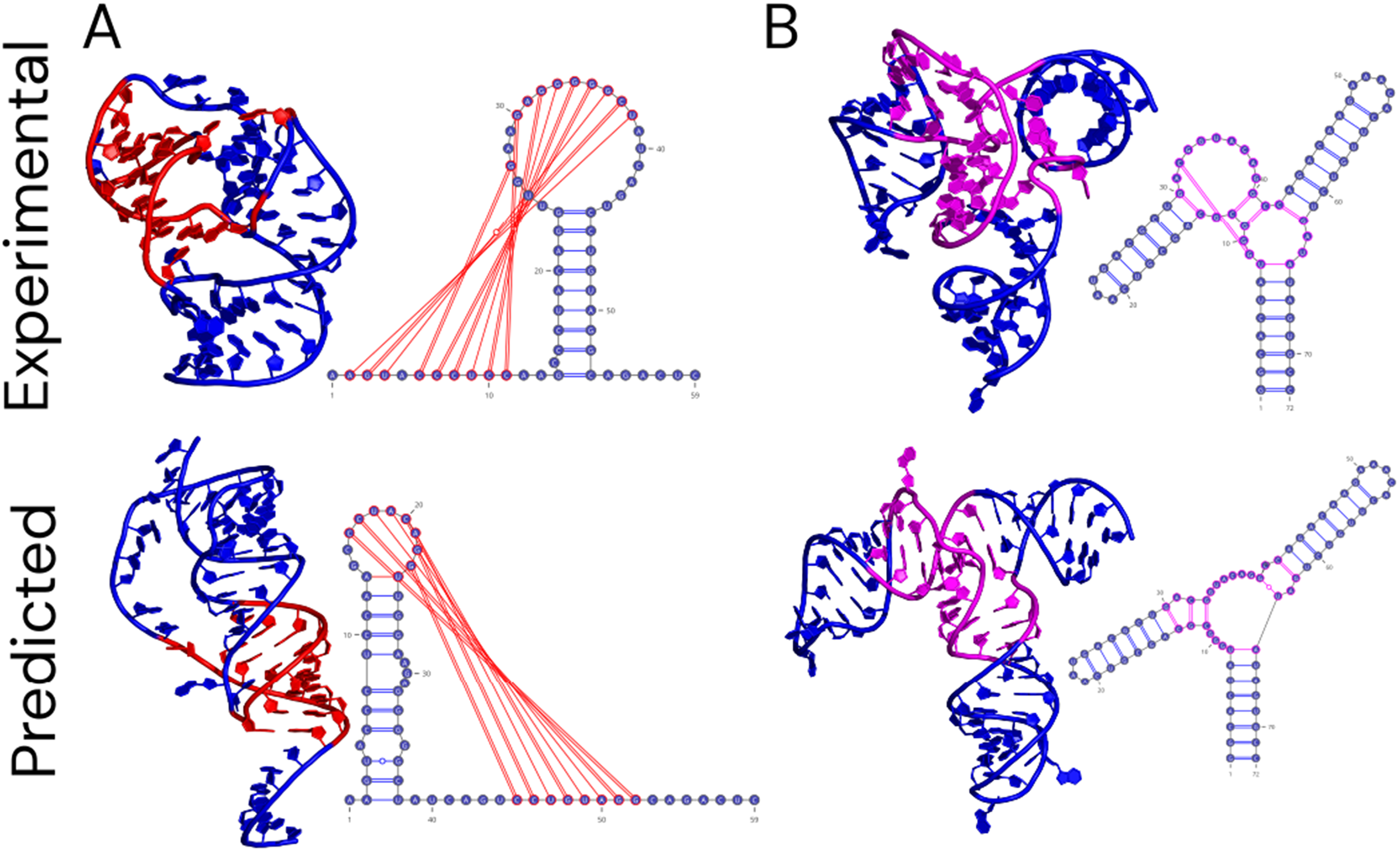
Errors in junction organization and tertiary contacts lead to incorrect global folds. (A) Target R1205 (xrRNA): while the correct pseudoknot topology was identified and enforced, the model failed to recover the native non-canonical base triples and specialized hairpin conformations, resulting in an incorrect global architecture. (B) Target R1209 (HIV-1 RRE stem–loop II): individual helical stems were captured with high accuracy, but incorrect coaxial stacking and helix orientation at the three-way junction core led to a misoriented, extended conformation. These examples illustrate the architectural barrier between local 2D fidelity and global 3D realism. Alt text: Experimental and predicted RNA structures are compared for two targets. In both cases, individual RNA structural elements are captured, but differences in junction organization, non-canonical base interactions, coaxial stacking, and helix orientation lead to incorrect global folds in the predicted structures.

A related failure mode is seen in R1209 (HIV-1 RRE stem–loop II; PDB ID: 9C2K), which illustrates how high local fidelity can coexist with global topological failure. Here, individual helical subdomains were captured with high accuracy—a result that mirrors the broad reliability of local elements observed in our hierarchical analysis (Figure 3B). However, the global fold remained incorrect due to the misorientation of these helices at the three-way junction core (Figure 4B). This confirms that the primary architectural challenge is not the prediction of helices themselves, but the recovery of the coaxial stacking and spatial connectivity (the components analyzed in Figure 3D) that define their relative arrangement. These cases illustrate how errors in junction organization can disrupt global three-dimensional architecture even when local secondary-structure elements are recovered.

#### 3.3.2 Mitigating failures through structural templates (M1209)

The prediction of target M1209 (the R1209 RNA bound to a chaperone Fab fragment; PDB ID: 9C2K) provides a vital contrast, showing how structural templates can bypass the architectural barriers that plague RNA-only modeling.

While the isolated R1209 monomer proved difficult due to its unconstrained junction, the protein–RNA complex was modeled with high accuracy (ICS and QS-scores ∼0.90). By leveraging homologous structures to map the conserved “GAAACAC” RNA binding motif, we were able to precisely dock the RNA onto the Fab scaffold (Figure 5). This case demonstrates that the availability of interfacial templates can mitigate the failures seen in unconstrained systems by providing the spatial priors that automated scoring functions currently fail to provide for *de novo* folds.

**Figure 5.**
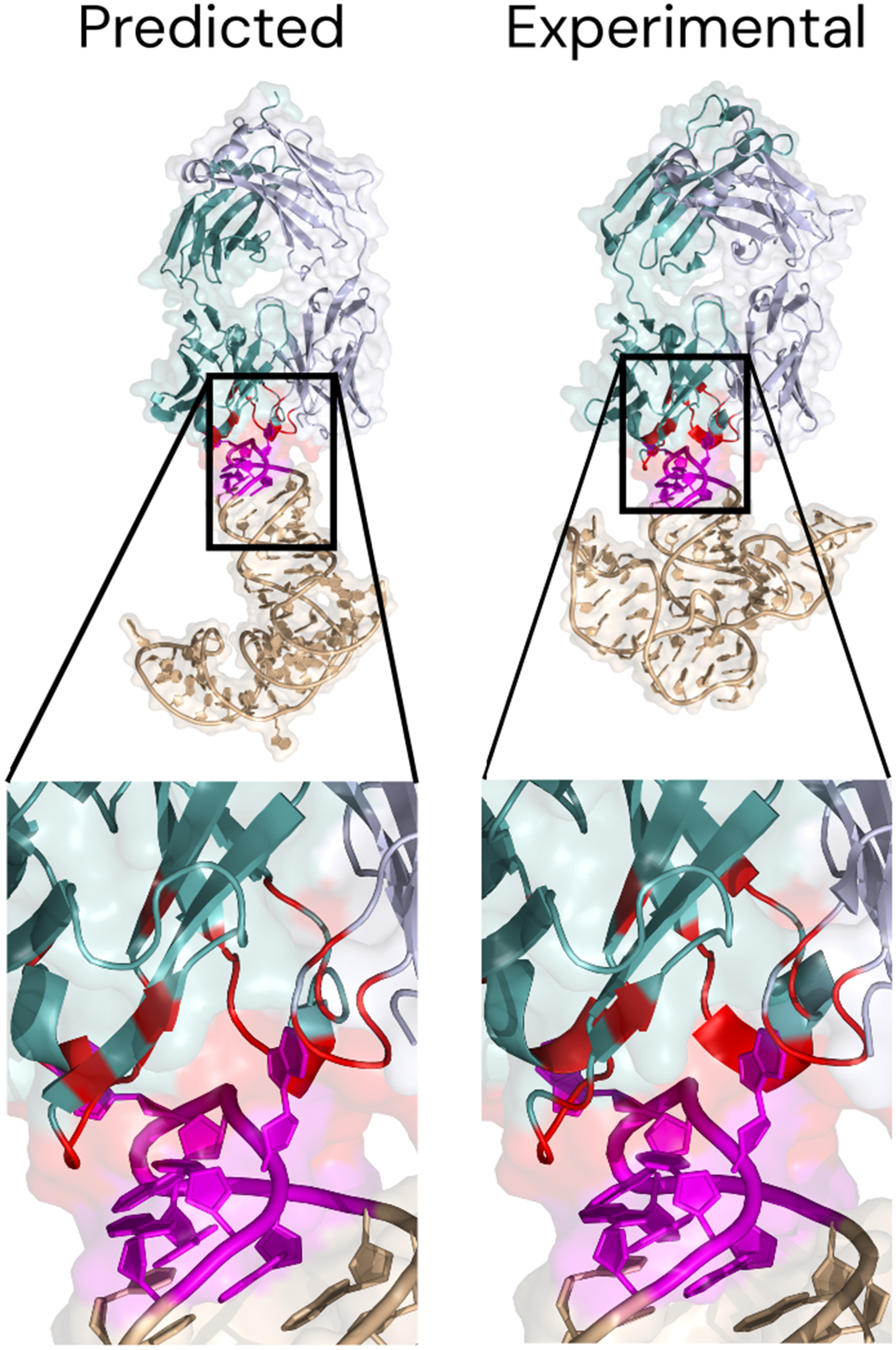
Accurate modeling of the RNA–protein interface in the M1209 complex. Experimental (left) and modeled (right) structures show close agreement at the protein– RNA interface (ICS and QS-scores ∼0.90). The use of homologous structural templates for the Fab chaperone and the conserved GAAACAC RNA binding motif (magenta) provided sufficient spatial constraints to bypass the junctional ambiguity observed in the isolated RNA monomer (R1209), demonstrating how interfacial templates can mitigate the failures seen in unconstrained systems. Alt text: Experimental and predicted structures of the M1209 RNA–protein complex are shown side by side, with enlarged views of the RNA–protein interface. The predicted interface closely resembles the experimental structure, including the conserved RNA binding region and associated protein contacts.

#### 3.3.3 Coarse-grained multimer organization despite limited atomic accuracy

The RNA–RNA multimer targets R1285 and R1283 exemplify the disconnect between benchmark ranking and structural realism identified in our analysis: high competitive rankings can be achieved despite limited absolute atomic precision.

For the OLE (ornate, large, and extremophilic) RNA homodimer (R1285o; PDB ID: 9MCW (Kretsch *et al*. 2025)), our pipeline successfully identified the correct dimeric stoichiometry and approximate binding mode, securing a top ranking in CASP16. However, the top-ranked model reached a global RMSD of approximately 11 Å (**Figure 6A**). This underscores the current frontier of the field: it is possible to recover the global organizational symmetry (the “coarse-grained” hypothesis) even while failing to reproduce the realistic interaction geometry and non-canonical edges that are systematically missed across multimeric targets (Figures 3E–F).

**Figure 6.**
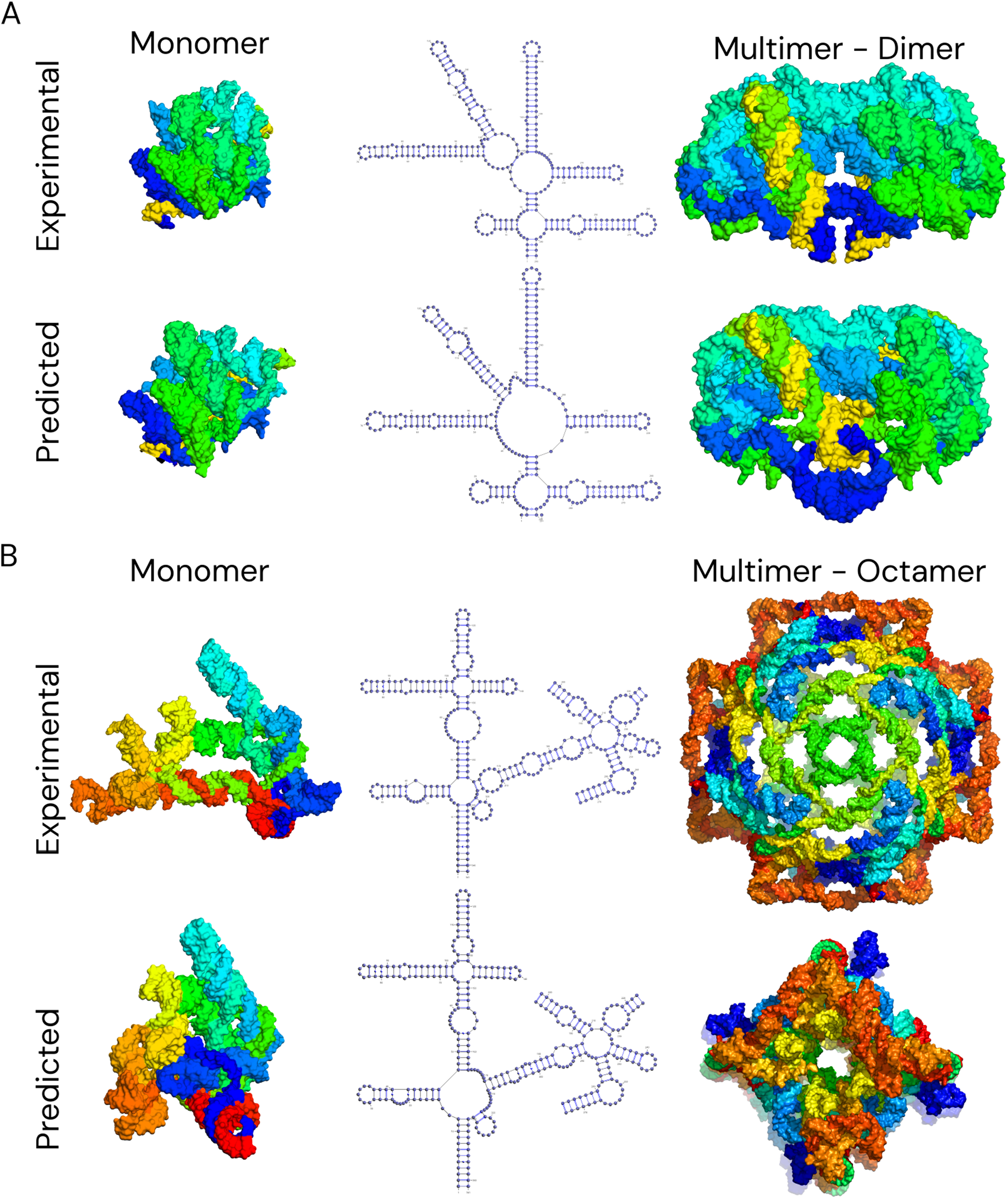
Coarse-grained recovery of RNA–RNA multimer assembly. **(A)** Target R1285 (OLE homodimer): the model correctly identified the dimeric stoichiometry and approximate arrangement of subunits, securing a top competitive ranking despite achieving low absolute atomic accuracy (global RMSD ∼11 Å). **(B)** Target R1283 v1/v3 (ROOL RNA nanostructure): while the monomeric building blocks were partially consistent with the reference architecture, the modeling pipeline failed to identify the native octameric symmetry, resulting in incomplete sub-assemblies. Secondary structure schematics are shown as VARNA diagrams. Alt text: Experimental and predicted RNA–RNA multimer assemblies are compared for two targets. For the dimeric target, the predicted monomer and multimer arrangements broadly reproduce the reference architecture. For the octameric target, the predicted monomers are partly consistent with the reference, but the assembled structure fails to reproduce the native octameric symmetry and remains incomplete.

A more complex failure occurred with R1283, a ROOL (rumen-originated, ornate, large) RNA nanostructure (Wang L *et al*. 2025). The challenge required predicting consistent architectures for monomeric (R1283v1; PDB ID: 9ISV), tetrameric (R1283v2; PDB ID: 9J3R) and octameric (R1283v3; PDB ID: 9J3T) assemblies. While individual monomeric subunits were partially consistent with the reference architecture, the pipeline failed to identify the correct multimeric assemblies, yielding only sub-assemblies. For the octamer (R1283v3o), AlphaFold 3 did not produce a physically plausible A8 model directly, and the submitted assembly was built by manually arranging A4 predictions into an A8 symmetry in PyMOL (Schrödinger and DeLano 2015), yielding only approximate sub-assemblies (**Figure 6B**). This confirms that the systematic exploration of higher-order assembly space remains a major challenge, as even small errors inherited from monomeric junctions propagate into large-scale shifts that preclude the formation of complex, symmetric interfaces.

To make the expert-guided component of the workflow more transparent, we added a target-level decision audit for RNA–RNA multimer targets in Supplementary Table S2. For each target or target group, the table summarizes the secondary-structure evidence, 3D and assembly sources, manual selection criteria, submitted-model rationale, post-CASP interpretation, and whether a more plausible retained candidate was available. This provides a target-level record of how individual modeling choices contributed to successes and failures in the final submissions.

## 4. Conclusions

This work provides a participant-level analysis of RNA structure prediction in CASP16, placing community-wide benchmark results in a direct structural context. By combining hierarchical performance statistics with detailed case studies, we distinguish between structural features that are currently more consistently captured by state-of-the-art pipelines and those that remain difficult to model in RNA prediction workflows. Our findings illustrate that prediction accuracy is not lost at a single point but instead declines progressively across the structural hierarchy. While current methods demonstrate reliability at the level of canonical base-pairing and local helical elements, this fidelity does not consistently translate into accurate tertiary organization or global architecture. Features that depend on non-canonical interactions, stacking geometry, junction organization, and long-range tertiary contacts are recovered less consistently and contribute to uncertainty in the resulting models.

The gradual loss of reliability across increasingly complex structural features provides a practical diagnostic for the field: it highlights where prediction shifts from relatively constrained local modeling toward more uncertain higher-order organization. This is particularly relevant for RNA–RNA multimers, where errors or uncertainties in monomer structure, non-canonical interactions, tertiary contacts, and interface geometry can propagate into assembly-level models. In the official CASP16 best-of-submitted-models analysis, our workflow achieved a top ranking in the RNA–RNA multimer category despite a more modest standing in the monomer assessment. This discrepancy suggests that, at the current state of the art, multimer success depends both on the ability to generate useful candidate models and on expert-guided assembly and model-selection decisions.

Ultimately, the limited recovery of specialized tertiary motifs, non-canonical interaction networks, and accurate tertiary or assembly-level geometries across the CASP16 dataset indicates that incremental improvements in sampling or refinement alone are unlikely to bridge the gap to atomic realism. The fact that multimer models currently operate as coarse-grained organizational hypotheses underscores the need for improvements at three linked levels: monomer structure accuracy, RNA–RNA interface modeling, and model selection under uncertainty. Future progress is unlikely to depend on structural data expansion alone. Given the limited number of experimentally determined RNA structures, improvements will also require stronger integration of physical and geometric principles into prediction and scoring, including better treatment of coaxial stacking, backbone geometry, ion- and solvent-mediated stabilization, and multimer assembly geometry. This is particularly important for stoichiometry, symmetry, and multimer assembly, where small upstream errors can lead to substantially different complex-level models.

## Acknowledgements

S.K., C.N., and S.P. acknowledge funding from the National Science Centre, Poland (2025/59/B/ST6/01643). We gratefully acknowledge Polish high-performance computing infrastructure PLGrid (HPC Centers: ACK Cyfronet AGH, WCSS) for providing computer facilities and support within computational grant nos. PLG/2025/017952 and PLG/2026/019105.

## Declaration of generative AI and AI-assisted technologies in the writing process

The authors employed ChatGPT (OpenAI), AI Studio (Google), Claude (Anthropic), and NotebookLM (Google) during the preparation of this manuscript and supplementary materials to facilitate drafting and to enhance the quality of the text. AI-assisted technologies were also used to assist in the preparation of Figure 1, including the generation and refinement of certain graphical elements and the overall visual presentation. For Figure 3, AI-assisted technologies were used to assist with the implementation and refinement of the underlying plotting code. The authors reviewed, verified, and, where necessary, revised the AI-assisted content, including the plotting code, and take full responsibility for the data, analyses, scientific content, visual design of the figures, and final publication.

## Data Availability Statement

The scripts used for feature extraction, scoring, bootstrap confidence-interval estimation, and figure generation are available at Zenodo: https://doi.org/10.5281/zenodo.21393731.

All models submitted to CASP16 are available for download from the official CASP website at: https://predictioncenter.org/download_area/CASP16/predictions/RNA/; https://predictioncenter.org/download_area/CASP16/predictions/oligo/; https://predictioncenter.org/download_area/CASP16/predictions/hybrid/.

All global and interface evaluation metrics used in this study were taken directly from the official CASP16 assessment and are available at: https://predictioncenter.org/casp16/results.cgi?tr_type=rna; https://predictioncenter.org/casp16/results.cgi?tr_type=rna_multi; https://predictioncenter.org/casp16/results.cgi?tr_type=hybrid.

## Conflict of Interest Statement

The authors declare that they have no competing interests.

## Supplementary Materials

Supplementary Methods S1. Detailed workflow for consensus-based RNA secondary structure prediction

Supplementary Methods S2. Detailed RNA 3D modeling workflow

Supplementary Methods S3. Official CASP16 metrics and post-CASP structural analyses

Supplementary Table S1: Hierarchical feature definitions and scoring framework

Supplementary Table S2: Target-level modeling decisions and post-CASP interpretation for RNA–RNA multimer targets

Supplementary Figure S1. Official CASP16 best-of-submitted-models performance rankings for the LCBio (G189) group

Supplementary Figure S2. Weighted heatmap of relative model performance for the LCBio (G189) group across CASP16 targets.

Supplementary Figure S3. Base interaction accuracy.

## Notes

https://doi.org/10.5281/zenodo.21393731

### Competing Interest Statement

The authors have declared no competing interest.

### Summary of Updates

The title and abstract are updated to reflect the reflect the intended contrast between relatively reliable local helical features and less reliable tertiary and assembly-level features.

