## Supplementary for "Hierarchical Breakdown of RNA Structure Prediction in CASP16: From Reliable Local Helices to Speculative Multimer Assembly"

for

### Supplementary Methods

**Supplementary Methods S1. Detailed workflow for consensus-based RNA secondary structure prediction**

#### S1.1 Initial secondary structure ensemble generation

For each RNA target, an initial ensemble of candidate secondary structures was generated using multiple independent prediction methods representing complementary methodological classes. Thermodynamic folding was performed using RNAfold from the ViennaRNA package and the Fold module of RNAstructure to obtain minimum free energy and suboptimal conformations. Probabilistic and machine learning–based approaches included CentroidFold , CONTRAfold , and RNAstructure MaxExpect. Pseudoknot-capable methods, ProbKnot , IPknot , and IPknot++, were additionally used to identify potential non-nested base-pairing interactions.

All tools in this ensemble were run with default parameters (no custom energy model, temperature, or window settings). RNAfold's minimum free energy structure was included in the consensus ensemble; the corresponding centroid structure was also generated but used only as a manual cross-check, not as an input to the automated consensus calculation. RNAstructure Fold generated a target-dependent number of suboptimal structures under default settings; only the lowest-energy structure was included in the consensus ensemble, since the suboptimal conformations are largely minor variants of the same fold and would otherwise dominate the ensemble by weight of numbers alone. Additionally, one structure each from the MaxExpect and ProbKnot programs of the RNAstructure package was included in the consensus ensemble. MaxExpect was included because it maximises expected base-pair accuracy rather than minimising free energy, and therefore represents a probabilistic view of the same thermodynamic model. ProbKnot was included because it derives base pairs from pairing probabilities without a nesting constraint, and so contributes pseudoknotted interactions that the thermodynamic and probabilistic methods cannot represent. For the dedicated pseudoknot predictors, both IPknot and IPknot++ were run, and the single structure with the greater number of predicted pseudoknots was included in the consensus ensemble; IPknot++ was used as the default choice when this comparison did not clearly favor one method. CentroidFold and CONTRAfold each contributed a single structure per target. Both were obtained using the CentroidFold implementation, which supports three energy schemes: McCaskill-BL, McCaskill-Turner and CONTRAfold. The CONTRAfold structure was generated under the CONTRAfold scheme within this implementation.

#### S1.2 Homology-based annotation and constrained folding

Sequence searches were performed using RNAcentral's own sequence search service . Match significance (E-value and bit score) was taken directly from RNAcentral's reported results; no additional custom threshold or manual judgment was applied on top of this. A hit was considered significant, and used, whenever RNAcentral reported one as such (e.g., a 134-nt HIV-1 RRE stem-loop matched a tRNA family, RF00005, at bit score 49.1 and E-value $2.4 \times 10^{-10}$; a 480-nt group II intron matched RF01998/RF02001 at bit scores 70.4 and 138.6, respectively). When a significant match to an Rfam family was identified by this case-by-case assessment, the corresponding secondary structure annotation was retrieved from R2DT. Reported homologs were not treated as experimentally validated structural templates; they were used only to guide consensus secondary-structure construction and manual inspection where family-level support was available. This homology-derived structure was subsequently used to guide constrained folding with RNAfold, using both soft and hard constraint regimes to generate additional structurally informed variants for inclusion in the candidate ensemble. When homologous templates linked to structural entries in PDBe were available, regions of the predicted secondary structures were manually adjusted to reflect conserved base-pairing patterns observed in the template structures, while allowing flexibility in unconserved regions.

##### S1.3 Local stability analyses for long RNAs

For longer RNAs, additional analyses were applied to improve robustness, using ScanFold and RNALfold.

ScanFold identifies locally stable structural motifs using randomized sequence backgrounds and Z-score analysis. ScanFold 2.0 was run with default parameters: a 120 nt window size, a 1 nt step size, a folding temperature of 37 °C, and mononucleotide shuffling to generate the randomized background. Base pairs passing a Z-score threshold of $\leq -2$, corresponding to highly stable structures, were retained as locally stable motifs.

RNALfold computes locally stable secondary structures with a maximal base-pair span using sliding window decomposition. RNALfold was run with the dangling-end model -d2 (the default) and a maximum base-pair span of 150 nt (the default), with Z-score filtering activated at a threshold of $\leq -2$, such that only locally optimal structures with a Z-score below this value were reported.

For shorter RNAs, these analyses were omitted, since the window sizes used by ScanFold 2.0 and RNALfold approach or exceed the full sequence length and the analyses therefore return no information beyond the global fold; in such cases, the consensus was derived exclusively from de novo thermodynamic, probabilistic, and pseudoknot-aware predictions.

##### S1.4 Covariance and multiple sequence alignment analysis

Multiple sequence alignments were generated using rMSA2 . These alignments were refined through covariance model–based searches. Covariance models were constructed with cmbuild and calibrated using cmcalibrate. They were then applied with cmsearch and cmalign to iteratively identify and align homologous sequences . Alignments were processed to remove poorly aligned regions. Columns corresponding to gaps in the reference sequence were also removed.

Secondary structures were inferred from the refined alignments using RNAalifold . Each candidate structure was evaluated against three criteria. First, covariation support was required, defined as at least four covarying base pairs across the alignment (MaxCovar $\geq 4$). Second, thermodynamic stability was assessed with AlifoldZ, with a threshold of Z-score $\leq -2.0$ . Third, RNAz was required to classify the region as a structured RNA, with an SVM RNA-class probability $\geq 0.80$ . AlifoldZ and RNAz were run independently. AlifoldZ tests whether the alignment folds more stably than shuffled alignments of the same composition. RNAz combines structure conservation and thermodynamic stability into a single classification.

#### S1.5 Consensus construction and restraint derivation

All predicted structures were converted to dot-bracket notation and retained for downstream consensus construction. At this stage, all methods were treated equally, and no weighting scheme was applied. Consensus base-pairing restraints were derived from base-pair occurrence frequencies across the ensemble of candidate secondary structures. Inclusion thresholds ranging from 0.0 to 1.0 were explored during consensus construction. This threshold reflects agreement across the independent methods contributing to the ensemble (e.g., RNAfold, IPknot, CentroidFold), not sampling frequency within any single method's suboptimal or stochastic output. A base pair at threshold 1.0 is therefore one recovered by every contributing method, regardless of how many suboptimal structures any individual method produced internally. For downstream three-dimensional modeling, two representative restraint sets were retained: a permissive set corresponding to a threshold of 0.5 and a stringent set corresponding to a threshold of 1.0. These thresholds were selected to balance structural inclusiveness against confidence and to bracket uncertainty during downstream three-dimensional modeling. The permissive set captured moderately supported interactions, whereas the stringent set retained only universally recurrent base pairs across all prediction methods.

#### S1.6 Manual curation and structural refinement

When mutually exclusive base-pairing configurations were observed, alternative pairing sets were preserved as separate structural hypotheses. Pseudoknot interactions that were not retained at the consensus level were manually reintroduced when supported by specific methods or by template-derived information.

Following consensus construction, candidate structures were manually inspected and curated using two-dimensional representations. Manual intervention was restricted to four defined stages: adjustment of template-informed regions, alignment cleaning, inspection of consensus structures, and targeted correction of implausible base-pairing patterns. Inspection criteria included helix continuity, absence of isolated base pairs, agreement with covariation signals, and preservation of known RNA architectural features such as junction topology and non-canonical interactions. Coaxial stacking arrangements were inferred from secondary-structure topology and geometric plausibility, considering helix continuity, junction connectivity, absence of intervening nucleotides, and consistency with covariation signals. These inferred stacking relationships were used as qualitative guidance during model evaluation and assembly and were not encoded as explicit base-pairing restraints.

Finalized consensus structures were converted into base-pairing restraints and used as primary inputs for downstream three-dimensional modeling.

**Supplementary Methods S2. Detailed RNA 3D modeling workflow**

#### S2.1. RNA 3D monomer modeling

Consensus-derived secondary structure restraints were used as inputs for three-dimensional RNA structure prediction. Candidate monomer models were generated using multiple complementary approaches: deep learning–based predictors, fragment-assembly methods, and simulation-based strategies. The methods used were DeepFoldRNA , RhoFold+ , NuFold , trRosettaNA , AlphaFold 3 , FARFAR2 , RNAComposer , Vfold2 , SimRNA , and BriQ .

Methods capable of incorporating secondary structure restraints were executed both with and without such restraints when applicable. For very long RNAs (>300 nt), computationally intensive methods (SimRNA, FARFAR2, DeepFoldRNA, RhoFold+ and Vfold2) were not applied. Across all methods, models generated under different conditions were treated as independent structural hypotheses and retained for downstream evaluation.

Deep learning–based predictors (DeepFoldRNA, RhoFold, NuFold, trRosettaNA, and AlphaFold 3) were run using sequence as the primary input. These methods internally generate multiple sequence alignments and derive structural restraints through neural network architectures. DeepFoldRNA produces up to six models per run, RhoFold+ generates a single model per prediction, NuFold generates up to four models per run, trRosettaNA generates up to five models per run, and AlphaFold 3 produces up to five models per run.

For most RNA monomer targets, AlphaFold 3 was run directly on the single-chain sequence. No additional stoichiometry or ion/cofactor screening was applied in these cases. Three monomer targets were treated differently. R1281 was screened with adenosine triphosphate (ATP), $Mg^{2+}$, a mixed $Mg^{2+}/Na^{+}$ condition, and nicotinamide adenine dinucleotide phosphate (NAP). R1286 was screened at three chain counts (1, 5, and 9 copies), with one 9-copy condition additionally including ATP. R1291 was screened with ATP, a mixed $K^{+}$/$Mg^{2+}$ condition, and NAP. In each case, AlphaFold 3 was run under these conditions to assess whether the presence of ions, cofactors, or additional chains improved prediction quality. Ions and cofactors were used only to aid structure prediction and were not retained in the final submitted coordinates. The complete stoichiometry and ion/cofactor screening procedure for RNA–RNA multimer targets is described in Supplementary Methods S2.3.

FARFAR2 simulations were performed for approximately one million Monte Carlo cycles per run, both with and without secondary structure restraints, generating up to five representative models per target . Vfold2 constructs RNA 3D structures through hierarchical assembly of helices, loops, and motifs derived from known structures . When Vfold2 pipeline is run without constraints, it internally generates alternative secondary structures using Vfold-2D, each leading to distinct 3D ensembles . Up to five representative models from these ensembles were retained. When secondary structure restraints were provided, a single ensemble was generated, from which up to five models were selected. RNAComposer was used as a fragment-based approach to generate models from provided secondary structure restraints, producing one model per restraint set .

SimRNA simulations employed a coarse-grained representation and were conducted both with and without secondary structure restraints using Replica Exchange Monte Carlo (REMC). For each RNA, eight independent trajectories were performed with different random seeds, each comprising ten replicas and 16 million iterations per run . Resulting conformations were clustered based on structural similarity using an RMSD cutoff corresponding to 10% of the sequence length, and representative low-energy structures from the top three clusters were selected following established protocols . BriQ , which relies on nucleobase-centric sampling and knowledge-based potentials, was executed with mandatory secondary structure restraints and produced a single model per simulation.

The total number of candidate models per target varied depending on sequence length, method applicability, and restraint conditions, but typically ranged from a few dozen to over one hundred across all methods.

#### S2.2. Refinement, filtering, and final model selection

Manual intervention in the 3D pipeline was applied at four defined stages: (i) manual PyMOL inspection during automated entanglement/clash filtering (below), (ii) manual comparison and superposition of candidate models to select for structural diversity in the final submitted set (below), (iii) for RNA–RNA multimer targets, manual visual screening of AlphaFold 3 predictions across candidate stoichiometries and ion/cofactor conditions, with manual duplication and repositioning of a physically plausible lower-order sub-assembly in

PyMOL when required (Supplementary Methods S2.3), and (iv) for RNA–protein complexes, manual docking guided by conserved interaction motifs and homologous interface templates, with local manual adjustment to resolve clashes and refine interface geometry (Supplementary Methods S2.4). These adjustments were limited to selecting, superposing, duplicating, repositioning, or locally resolving clashes in candidate models; no unsupported sequence register changes or de novo redesign of RNA topology were introduced during final model selection.

Throughout this pipeline, a model was classified as having non-physical geometry if it exhibited any of the following: topological chain entanglement (linked or threaded backbone paths, identified by RNAspider ); severe steric clashes between non-bonded atoms; spatial self-overlap within a single chain, or overlap between distinct chains occupying the same three-dimensional region; or helical-like backbone conformations lacking the base-pairing interactions required to support a helix. This definition was applied consistently across both the automated and manual filtering steps described below.

Across all methods, multiple candidate models were generated per target and retained for downstream evaluation. Clustering and energy-based filtering were applied within individual methods, while cross-method comparison focused on structural consistency and agreement with input secondary structure restraints.

All retained candidate models underwent refinement with QRNAS using default settings for 5000 steps to improve stereochemistry and reduce structural clashes. Models were then filtered using a combination of automated and manual criteria.

Automated filtering included analysis with RNAspider to identify structural entanglements. Models exhibiting topological entanglements were removed unless no alternative model without entanglement was available. Manual inspection was performed in PyMOL to assess structural plausibility. Models were discarded if they contained severe steric clashes, overlapping chains occupying the same spatial region, self-overlapping segments, or helical-like conformations lacking proper base pairing.

Final submitted models were selected through manual inspection with emphasis on both structural plausibility and conformational diversity. Candidate models were visually compared and superposed to identify structurally similar groups, approximating clustering without formal RMSD-based procedures. Assessment of structural similarity and diversity was performed by expert inspection, guided by differences in global fold topology (e.g., helix arrangement and relative orientation), junction configuration, and long-range contacts; models exhibiting the same overall fold were considered redundant despite minor local variations. Representative models were then selected to maximize coverage of distinct conformational states while avoiding redundancy.

#### S2.3. RNA–RNA multimer assembly procedure

Assembly of RNA–RNA multimers was performed as an expert-guided, heuristic procedure rather than an automated docking or scoring pipeline.

For RNA–RNA multimer targets, candidate higher-order assemblies were generated by evaluating alternative stoichiometries and assembling monomer models into putative complexes. Stoichiometry was inferred empirically by running AlphaFold 3 directly at multiple candidate chain counts, typically spanning 1 to 8 copies and up to 12 for the largest assemblies. Chain counts were combined with ion and cofactor conditions on a target-specific basis rather than exhaustively; the conditions evaluated for each target are listed in Supplementary Table S2. The tested conditions comprised a baseline of RNA alone, $Mg^{2+}$, ATP, NAP, palmitic acid (PLM), a mixed $Mg^{2+}/Na^{+}$ condition, a ten-ion cocktail ($Mg^{2+}$, $Na^{+}$, $K^{+}$, $Co^{2+}$, $Zn^{2+}$, $Ca^{2+}$, $Mn^{2+}$, $Fe^{2+}$, $Cu^{2+}$ and $Cl^{-}$), and combinations of these. Ions and cofactors were included to assess whether their presence reduced chain entanglement and crossing and improved prediction quality. Configurations that produced severe steric clashes, chain entanglement, or overlapping chains were discarded on visual inspection. No automated scoring function was used to rank candidate stoichiometries or assemblies against one another.

AlphaFold 3 could not always produce a physically plausible prediction at the full target stoichiometry, for example due to entanglement at higher copy numbers. In such cases, a lower-order sub-assembly that AlphaFold 3 predicted natively and without entanglement, such as a physically plausible tetramer, was built independently. This sub-assembly was then manually duplicated and repositioned in PyMOL to approximate the required higher-order symmetry. Because no validated quantitative discriminator was available to rank alternative manual placements, the final choice relied on expert visual judgment. This is a specific source of uncertainty in the resulting higher-order assemblies.

Symmetry relationships were visually approximated during model construction. When structurally similar assemblies were available in the Protein Data Bank , monomer models were crudely superposed onto template architectures in PyMOL to guide complex assembly. Known interactions described in the literature were preserved where relevant.

Monomer structures were treated as rigid or semi-rigid bodies and assembled through manual docking in PyMOL. Interface evaluation was based on visual inspection, focusing on steric compatibility, continuity of helices across interfaces, and overall geometric plausibility. No automated scoring or global optimization procedure was applied.

Target-specific detail on the exact chain counts and ion/cofactor conditions evaluated, which sub-assembly (if any) required manual duplication, and whether a better-scoring alternative existed among the retained candidates but was not selected as the primary submitted model, is provided for each RNA–RNA multimer target in Supplementary Table S2.

##### S2.4. RNA–protein complex assembly procedure

Assembly of RNA–protein complexes were assembled as an expert-guided, heuristic procedure rather than an automated docking pipeline.

For RNA–protein complexes, an explicit interface prediction step preceded assembly. Sequence similarity searches and literature-based curation were used to identify conserved interaction motifs and homologous interfaces. These features were translated into qualitative spatial restraints to guide assembly of independently modeled RNA and protein components.

Complexes were assembled manually in PyMOL and refined through local adjustments to resolve steric clashes and improve interface geometry. No exhaustive docking or energy-based ranking was performed.

#### Supplementary Methods S3. Official CASP16 metrics and post-CASP structural analyses

##### S3.1. Weighted relative performance

To construct the global relative performance heatmap, the predicted quality for the LCBio group was mapped against the community distribution using standard scores (Z-scores). All global and interface evaluation metrics used in this study were taken directly from the official CASP16 assessment (https://predictioncenter.org/download_area/CASP16/) and were not recomputed . For each CASP16 target and each metric independently, the community mean and standard deviation were computed from the full set of submitted predictors. The LCBio model deviation was then expressed as a Z-score, defined as the number of standard deviations from the community mean. To account for scoring directionality, Z-scores were inverted for metrics where lower values correspond to better performance (e.g., RMSD, clash score). The resulting matrices were transposed to arrange evaluation metrics along one axis and targets along the other.

To summarize performance across metrics, a composite score (Zmonomer or Zmultimer) was computed and appended alongside the individual metrics, following the official CASP evaluation scheme . For monomeric predictions, this score was calculated as a weighted linear combination of TM-score (0.3), GDT_TS (0.3), and lDDT (0.4). For RNA–RNA multimeric complexes, the score combined the monomeric component (0.3) with interface-specific metrics (ICS, IPS, and ilDDT; total weight 0.7). To reduce the influence of extreme outliers on visualization, negative Z-scores were capped at −2.0. The final matrices were rendered using a diverging color map centered at zero (Z = 0), indicating performance above (blue) or below (red) the community mean for each target and metric.

#### S3.2. Feature-level structural analysis across the RNA structural hierarchy

All structural features used in the analysis were derived from reference and predicted RNA structures using DSSR , which provided a standardized annotation of base pairs, stacking interactions, secondary structure elements, junctions, and tertiary motifs. All global fold and interface metrics used in this study were taken directly from the official CASP16 assessment and were not recomputed. Feature-level analyses were performed on the models submitted by the LCBio group in CASP16 to quantify prediction accuracy across hierarchical levels of RNA structural organization.

For each target, a single representative model was selected based on the highest composite score (Zmonomer or Zmultimer) and compared against the corresponding reference structure after sequence alignment. This model was selected as the representative to align feature-level analysis with the official CASP ranking criteria, ensuring that the most competitive submission per target was used for detailed evaluation. Accuracy for interaction-based features was quantified using the F1-score based on counts of true positives (TP), false positives (FP), and false negatives (FN). For higher-level structural categories, counts were aggregated across contributing sub-features before F1-score calculation. A summary of the hierarchical feature definitions and corresponding scoring procedures is provided in Supplementary Table S1.

Watson–Crick base pairs were evaluated using strict matching, where a predicted pair was considered correct only if both residues corresponded exactly to the reference pair after sequence mapping.

Non-Watson–Crick base pairs were evaluated using an explicit 6-point rubric, split as: 2 points for identity (confirming the pair occurs between the same two residues, after sequence-alignment mapping, in both reference and model), 2 points for matching cis/trans orientation, and 1 point each for the two Leontis–Westhof interacting edges (one per partner base). A reference pair absent from the model at the mapped position contributes its full 6 points to false negatives; a predicted pair with no corresponding reference position contributes its full 6 points to false positives. When a pair is identified in both but with a mismatched orientation and/or edge classification, only the matching components count toward true positives, and each mismatched component contributes to both a false positive and a false negative — so a partially-correct interaction is penalized on the specific component that is wrong rather than being scored all-or-nothing. These per-component counts are summed across all interactions in a target/model and combined into the reported F1-score.

Base stacking interactions were evaluated as pairwise contacts derived from contiguous stacking segments, using the same 6-point structure as non-Watson–Crick pairs: 2 points for identity, 2 points for inter-planar distance matching the reference within 0.5 Å, and 2 points for angular deviation between base normals within 15° (1 point is awarded for each threshold independently satisfied where applicable). Deviations from these thresholds contributed to partial penalties on the same true/false positive and negative basis described above. Residues annotated as non-stacking in the reference were evaluated separately to penalize overprediction of stacking interactions.

Secondary structure elements, including helices, stems, hairpins, bulges, internal loops, and single-stranded segments, were evaluated using a motif-based matching procedure. Reference and predicted motifs were first aligned based on residue overlap, requiring more than 50% overlap to establish correspondence. Once matched, accuracy was computed at the level of constituent units, defined as base pairs for helices and stems, and residues for loop and single-stranded elements. Unmatched motifs contributed fully to false positive or false negative counts.

Junction geometry was evaluated using a topology-gated framework. A predicted junction was first required to match the reference in terms of junction type, defined by the number of outgoing helices. Only junctions satisfying this condition were further evaluated. Four components were then assessed: (i) connectivity, defined by the ordering of helices; (ii) recovery of terminal base pairs at helix ends; (iii) preservation of linker residues as unpaired; and (iv) identification of coaxial stacking relationships between helices. Each component contributed proportionally to the total score, allowing partial credit for partially correct junctions. Junctions were additionally stratified by topology to evaluate performance across different branching complexities.

Global fold accuracy was quantified using the TM-score provided for each model. This metric was not recomputed but directly incorporated into the hierarchical analysis as the measure of global structural agreement.

Tertiary motifs included A-minor interactions, ribose zippers, K-turns, kissing loops, pseudoknots, and base multiplets, were evaluated using the same motif-matching framework applied to secondary structure elements. Motifs were matched based on residue overlap and scored at the level of constituent residues or interactions.

For RNA–RNA multimer targets, interface accuracy was defined as recovery of inter-chain base pairs and does not incorporate global interface metrics (e.g., ICS, IPS, ilDDT).

#### S3.3 Cluster bootstrap confidence intervals

Error bars were obtained by a cluster (block) bootstrap with 2,000 resamples per estimate, taking the 2.5th and 97.5th percentiles of the resulting distribution as a 95% confidence interval. The ten scored RNA–RNA multimer targets are not independent observations: several represent different stoichiometries or conformations of the same underlying RNA. These were grouped into seven resampling units: GOLLD (R1250o, R1254o); three ROOL units reflecting three distinct underlying RNAs, namely ROOL-Lactobacillus (R1252o), ROOL-env209 (R1253v1o, R1253v2o), and ROOL-Enterococcus (R1283v2o, R1283v3o); and three further targets from unrelated RNA classes, each treated as its own unit, namely OLE (R1285o), the 6-helix bundle (R1281o), and HYER1 (R1290o). At each bootstrap iteration, whole units rather than individual targets were resampled with replacement from this set of seven. Monomeric targets, which do not share this family structure, were resampled individually. For quantities reported as mean F1-scores across targets, each iteration resampled per-target values and recomputed the mean. For quantities reported as percentage recovery of reference interactions, each iteration instead resampled per-target raw interaction counts and recomputed the aggregate percentage from the pooled counts, since these percentages are ratios and cannot be correctly re-estimated by resampling the percentages themselves.

#### Supplementary Table S1: Hierarchical feature definitions and scoring framework

| Structural feature | Description | Matching criterion | Scoring approach |
|---|---|---|---|
| Watson–Crick base pairs | Canonical base pairs | Exact correspondence of both residues after sequence alignment | Correct pairs counted as true positives; mismatches contribute to false positives and false negatives; F1-score computed from TP/FP/FN |
| Secondary structure elements | Helices, stems, hairpins, bulges, internal loops, single-stranded regions | ≥50% overlap between mapped reference and predicted elements | Overlapping units contribute to true positives; non-overlapping regions contribute to false positives and false negatives; F1-score computed from aggregated counts |
| Junction geometry | Multi-helix junctions (e.g., 3-way, 4-way) | Junction type (number of helices) must match; otherwise counted as incorrect | Correct junctions evaluated by helix connectivity, terminal base-pair recovery, linker residues remaining unpaired, and coaxial stacking; partial credit aggregated into TP/FP/FN and converted to F1 |
| Non-Watson–Crick interactions | Non-canonical base pairs classified by Leontis–Westhof geometry | Same pair of residues must be identified | Scoring decomposed into orientation (cis/trans) and interacting edges; correct components contribute partial credit; totals accumulated into TP/FP/FN and converted to F1 |
| Base stacking | Stacking interactions between adjacent bases | Same pair of residues must be identified | Scoring based on geometric agreement (distance and relative orientation); correct components contribute partial credit; totals aggregated into TP/FP/FN and converted to F1 |
| Global fold | Overall structural agreement | Direct comparison to reference | TM-score reported from CASP assessment used directly |
| Tertiary motifs | A-minor interactions, ribose zippers, k-turns, kissing loops, pseudoknots, multiplets | ≥50% overlap between mapped reference and predicted motifs | Overlapping units contribute to true positives; unmatched regions contribute to false positives and false negatives; F1-score computed |
| RNA–RNA interface | Inter-chain base-pairing interactions | Exact correspondence of interacting residues | Correct pairs contribute to true positives; incorrect or missing pairs contribute to false positives and false negatives; F1-score computed |

**Supplementary Table S2. Target-level modeling decisions and post-CASP interpretation for RNA–RNA multimer targets.** If a specific tool or decision could not be uniquely assigned as responsible for the final outcome, this is indicated explicitly. Post-CASP interpretations are based on comparison with the released experimental structures and are intended to document likely sources of success or failure, not to imply that these factors were known during blind prediction.

| RNA type | Target (Length, Stoich) | 2D evidence | 3D/ assembly source | Manual decision and submitted rationale | Post-CASP interpretation[#] | Better retained model available? |
|---|---|---|---|---|---|---|
| GOLLD | R0250o (744 nt, UNK) | RNACentral searches identified homologous sequences associated with Rfam family RF02032. A consensus 2D structure was generated using: RNAfold, RNAstructure (Fold, MaxExpect, ProbKnot), IPknot, CentroidFold (McCaskill-BL/Turner, ContraFold), RNAalifold with default parameters. The RNAalifold predictions were evaluated with AlifoldZ (Z-score ≤ −2.0) and RNAz (score ≥ 0.80) for inclusion. The 2D structure was used as guidance for visual inspection of 3D models. | AlphaFold 3 (AF3) was run systematically from A1 to A6 stoichiometries. | Four out of five models submitted were monomers while one was a dimer based on visual screening. | All submitted models had wrong stoichiometry. | No. Other retained models were either similar to the submitted models or had non-physical geometries, including chain overlap, chain crossings, or entanglements. |
| GOLLD | R1250o (744 nt, A6) | Predictions from phase 0 were reused in phase 1. | Predictions from phase 0 were reused in phase 1. | No models were submitted. | Not applicable. | No. All 3D models from A6 stoichiometry had non-physical geometries, including chain overlap, chain crossings, or entanglements. |

| | | | | | | |
|---|---|---|---|---|---|---|
| GOLLD | R0254o (413 nt , UNK) | RNACentral searches identified homologous sequences associated with Rfam family RF02032. A consensus 2D structure was generated using RNAfold, RNAstructure (Fold, MaxExpect, ProbKnot), IPknot, CentroidFold (McCaskill-BL/Turner, ContraFold), RNAalifold with default parameters. The RNAalifold predictions were evaluated with AlifoldZ (Z-score ≤ −2.0) and RNAz (score ≥ 0.80) for inclusion. The 2D structure was used as guidance for visual inspection of 3D models. | AF3 was run systematically from A1 to A12 stoichiometries with and without $Mg^{2+}$ ions. Additionally, monomers were built with DeepFoldRNA, Nufold, RhoFold, trRosettaRNA and RNAComposer. All monomer models were discarded based on visual inspection. | All submitted models were hexamers picked from AF3 runs without $Mg^{2+}$ ions. | All submitted models had wrong stoichiometry. | No. Other retained models were either similar to the submitted models or had non-physical geometries, including chain overlap, chain crossings, or entanglements. |
| GOLLD | R1254o (413 nt, A14) | Predictions from phase 0 were reused in phase 1. | Predictions from phase 0 were reused in phase 1. | The AF3 models from heptamer predictions were manually arranged into A14 stoichiometry in PyMOL. | All five models ranked poorly and inconsistently across metrics. | No. No better models were available because the pool was limited to five hand-assembled models. |

| | | | | | | |
|---|---|---|---|---|---|---|
| OLE | R0285o (577 nt, UNK) | RNACentral searches identified homologous sequences associated with Rfam family RF01071. A consensus 2D structure was generated using RNAfold, RNAstructure (Fold, MaxExpect, ProbKnot), IPknot, CentroidFold (McCaskill-BL/Turner, ContraFold), RNAalifold with default parameters. The RNAalifold predictions were evaluated with AlifoldZ (Z-score ≤ −2.0) and RNAz (score ≥ 0.80) for inclusion. The 2D structure was used as guidance for visual inspection of 3D models. | AF3 was run systematically from A1 to A8 stoichiometry. All simulations were run in presence of co-factor ATP and $Mg^{2+}$ ions to reduce entanglements and chain crossings and to improve scores. | Dimer (model one) and trimer (model two) were the two most plausible of the 8 stoichiometry hypotheses screened based on visual inspection. The remaining submitted models were monomers. | The correct stoichiometry (dimer) was identified in the primary model. It was also the best-ranked model submitted by any group. | No. Other retained models were either similar to the submitted models or had non-physical geometries, including chain overlap, chain crossings, or entanglements. |
| OLE | R1285o (577 nt, A2) | Predictions from phase 0 were reused in phase 1. | AF3 was run with A2 stoichiometry. The simulations were performed under six conditions with: baseline, $Mg^{2+}$, combination of ten ions, ATP, ions $Mg^{2+}$ + $Na^{+}$ and NAP. The baseline condition included RNA only. The combination of 10 ions included $Mg^{2+}$, $Na^{+}$, $K^{+}$, $Co^{2+}$, $Zn^{2+}$, $Ca^{2+}$, $Mn^{2+}$, $Fe^{2+}$, $Cu^{2+}$, and $Cl^{-}$. | All models were selected based on visual inspection. Only two of the six screened conditions (Combination of ten ions, Mg-only) contributed to the submission. The order of models was determined based on visual confidence. All submitted models were manually adjusted by hand in PyMOL. | Among the submitted models, model four was the best. It was also the best-ranked model submitted by any group. | No. Other retained models were either similar to the submitted models or had non-physical geometries, including chain overlap, chain crossings, or entanglements. |

| | | | | | | |
|---|---|---|---|---|---|---|
| ROOL | R1052o (520 nt, UNK) | A consensus 2D structure was generated using RNAfold, RNAstructure (Fold, MaxExpect, ProbKnot), IPknot, CentroidFold (McCaskill-BL/Turner, ContraFold), RNAalifold with default parameters. The RNAalifold predictions were evaluated with AlifoldZ (Z-score ≤ −2.0) and RNAz (score ≥ 0.80) for inclusion. The 2D structure was used as guidance for visual inspection of 3D models. | AF3 was run with A2, A3 and A6 stoichiometries. Two additional runs were performed for A6 with $Mg^{2+}$ and ATP/NAP. | No models were submitted. | Not applicable. | Not applicable. |
| ROOL | R1252o (520 nt, A6) | Predictions from phase 0 were reused in phase 1. | Predictions from phase 0 were reused in phase 1. | Models one and two were from A6 AF3 runs with $Mg^{2+}$ and ATP. Models three and four were hand-built in PyMOL from the A3 AF3 models while model five was hand-built using the A2 AF3 models. | Hand-built A6 model (model three) ranked three among all models submitted by any group. All other models ranked poorly. | No. Other retained models were either similar to the submitted models or had non-physical geometries, including chain overlap, chain crossings, or entanglements. |
| ROOL | R0253o (574 nt, UNK) | A consensus 2D structure was generated using RNAfold, RNAstructure (Fold, MaxExpect, ProbKnot), IPknot, CentroidFold (McCaskill-BL/Turner, ContraFold), RNAalifold with default parameters. The RNAalifold predictions were evaluated with AlifoldZ (Z-score ≤ −2.0) and RNAz (score ≥ 0.80) for inclusion. The 2D structure was used as guidance for visual inspection of 3D models. | AF3 was run with A8 stoichiometry under 4 conditions: $Mg^{2+}$, Palmitic Acid (PLM), NAP, $Mg^{2+}$ + ATP + NAP. | No models were submitted. | Not applicable. | Not applicable. |

| | | | | | | |
|---|---|---|---|---|---|---|
| ROOL | R1253v1o (574 nt, A8, Conf 1) | Predictions from phase 0 were reused in phase 1. | Predictions from phase 0 were reused in phase 1. | Model 1 was from AF3 run with $Mg^{2+}$ ions, models 2-4 were from runs with PLM and model 5 from $Mg^{2+}$ + ATP + NAP run. | The models five and one were ranked two and three among all models submitted by any group. | Yes. Models similar to the two higher-ranked submissions were present in the retained pool but were not selected. |
| ROOL | R1253v2o (574 nt, A8, Conf 2) | Predictions from phase 0 were reused in phase 1. | Predictions from phase 0 were reused in phase 1. | Same set of models were used as conformation 1. | The models five and one were ranked seven and eight among all models submitted by any group. | Yes. Models similar to the two higher-ranked submissions were present in the retained pool but were not selected. |
| ROOL | R0283o (580 nt, UNK) | A consensus 2D structure was generated using RNAfold, RNAstructure (Fold, MaxExpect, ProbKnot), IPknot, CentroidFold (McCaskill-BL/Turner, ContraFold), RNAalifold with default parameters. The RNAalifold predictions were evaluated with AlifoldZ (Z-score ≤ −2.0) and RNAz (score ≥ 0.80) for inclusion.<br>The 2D structure was used as guidance for visual inspection of 3D models. | AF3 predictions were run with stoichiometries A1–A8, not uniformly. A1 and A2 were each simulated under 3 conditions: baseline, with ATP, with NAP. A4 was simulated under 2 conditions: with ATP, with NAP. A3, A5, A6, A7, and A8 were simulated only once, with ATP. | The submitted models were individual chains extracted from runs with A4, A6 and A8 stoichiometries. | All submitted models had wrong stoichiometry. | Yes. There were better models in A4 stoichiometry predictions and were submitted in phase 2. |
| ROOL | R1283v2o (580 nt, A4) | Predictions from phase 0 were reused in phase 1. | Predictions from phase 0 were reused in phase 1. | Models were submitted based on visual inspection. All models were from A4 runs, first four from runs with ATP while the fifth one from runs with NAP. | Model one ranked three among all models submitted by any group. | No. Other retained models were either similar to the submitted models or had non-physical geometries, including chain overlap, chain crossings, or entanglements. |

| | | | | | | |
|---|---|---|---|---|---|---|
| ROOL | R1283v3o (580 nt, A8) | Predictions from phase 0 were reused in phase 1. | Predictions from phase 0 were reused in phase 1. | Models were submitted based on visual inspection. Most of the models generated by AF3 were non-physical. The AF3 models from A4 were adjusted by hand into A8 in PyMOL. | Models 1-3 ranked 1-3 among all models submitted by any group. However, these models are far from the reference structure with RMSDs in the 99.1–107.6 Å range. | No. Other retained models were either similar to the submitted models or had non-physical geometries, including chain overlap, chain crossings, or entanglements. |
| 6-helix bundle | R0281o (718 nt, UNK) | A consensus 2D structure was generated using RNAfold, RNAstructure (Fold, MaxExpect, ProbKnot), IPknot, CentroidFold (McCaskill-BL/Turner, ContraFold), RNAalifold with default parameters. The RNAalifold predictions were evaluated with AlifoldZ (Z-score ≤ −2.0) and RNAz (score ≥ 0.80) for inclusion. The 2D structure was used as guidance for visual inspection of 3D models. | AF3 was run with A2 and A3 stoichiometries. Two additional runs were performed for A2 with ATP and NAP. | No models were submitted. | Not applicable. | No. All 3D models had non-physical geometries, including chain overlap, chain crossings, or entanglements. |
| 6-helix bundle | R1281o (718 nt, A2) | Predictions from phase 0 were reused in phase 1. | Using the template PDBs 7PTK, 7PTL, 7PTQ and 7PTS homology models were built using ModeRNA. | The homology models were were adjusted by hand to A2 stoichiometry in PyMOL. | All five models ranked poorly and inconsistently across metrics. | No. No better models were available because the pool was limited to five hand-assembled models. |

| | | | | | | |
|---|---|---|---|---|---|---|
| HYER1 | R0290o (627 nt, UNK) | A consensus 2D structure was generated using RNAfold, RNAstructure (Fold, MaxExpect, ProbKnot), IPknot, CentroidFold (McCaskill-BL/Turner, ContraFold), RNAalifold with default parameters. The RNAalifold predictions were evaluated with AlifoldZ (Z-score ≤ −2.0) and RNAz (score ≥ 0.80) for inclusion. The 2D structure was used as guidance for visual inspection of 3D models. | No 3D predictions were performed in phase 1. | No models were submitted. | Not applicable. | Not applicable. |
| HYER1 | R1290o (627 nt, A2) | Predictions from phase 0 were reused in phase 1. | AF3 was run with A2 stoichiometry. The simulations were performed under six conditions with: baseline, $Mg^{2+}$, combination of ten ions, ATP, ions $Mg^{2+}$ + $Na^{+}$ and NAP. The baseline condition included RNA only. The combination of 10 ions included $Mg^{2+}$, $Na^{+}$, $K^{+}$, $Co^{2+}$, $Zn^{2+}$, $Ca^{2+}$, $Mn^{2+}$, $Fe^{2+}$, $Cu^{2+}$, and $Cl^{-}$. | All models were selected based on visual inspection. Only two of the six screened conditions (Combination of ten ions, $Mg^{2+}$ + $Na^{+}$) contributed to the submission. The order of models was determined based on visual confidence. All submitted models were adjusted by hand in PyMOL. | All five models ranked poorly and inconsistently across metrics. | No. Other retained models were either similar to the submitted models or had non-physical geometries, including chain overlap, chain crossings, or entanglements. |

[#] Rankings refer to the official CASP16 best-of-submitted-models assessment unless otherwise stated.

#### Supplementary Figures

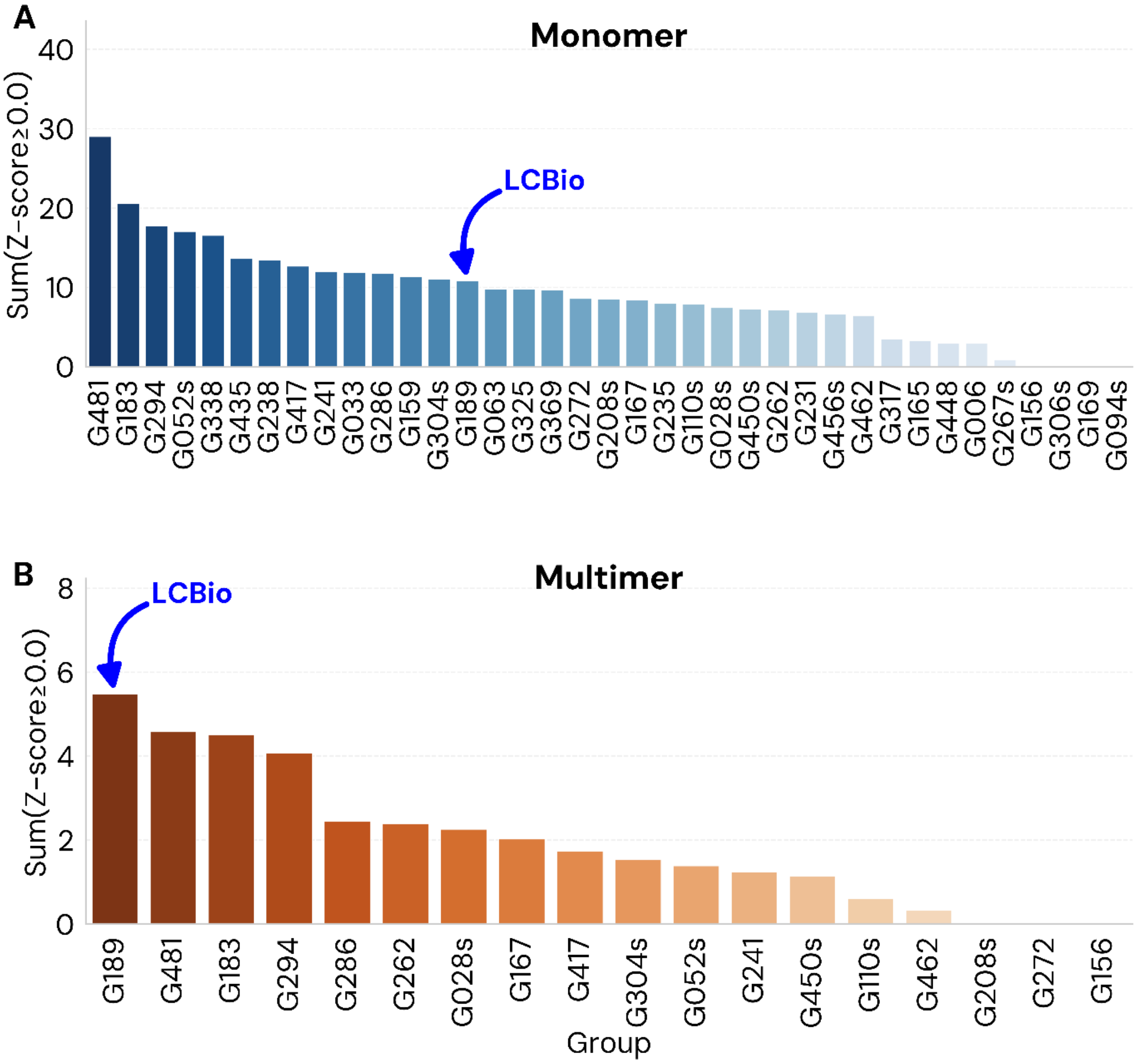


**Supplementary Figure S1. Official CASP16 best-of-submitted-models performance rankings for the LCBio (G189) group.** (A) Ranking across monomeric RNA targets. (B) Ranking across RNA–RNA multimer targets. For each panel, groups are ranked by their cumulative performance across all targets in the respective category, expressed as the sum of positive Z-scores (Sum of $Z_{monomer}$ or $Z_{multimer} \geq 0.0$). This metric, taken from the official CASP16 assessment , summarizes overall structural accuracy and interface quality. Columns represent individual participating groups (e.g., G481, G183), with the LCBio group (G189) highlighted by a blue arrow and text label. Groups whose identifier ends in ’s’ denote server submissions. The color gradients (blue for monomers and orange for multimers) reflect relative ranking within each category.

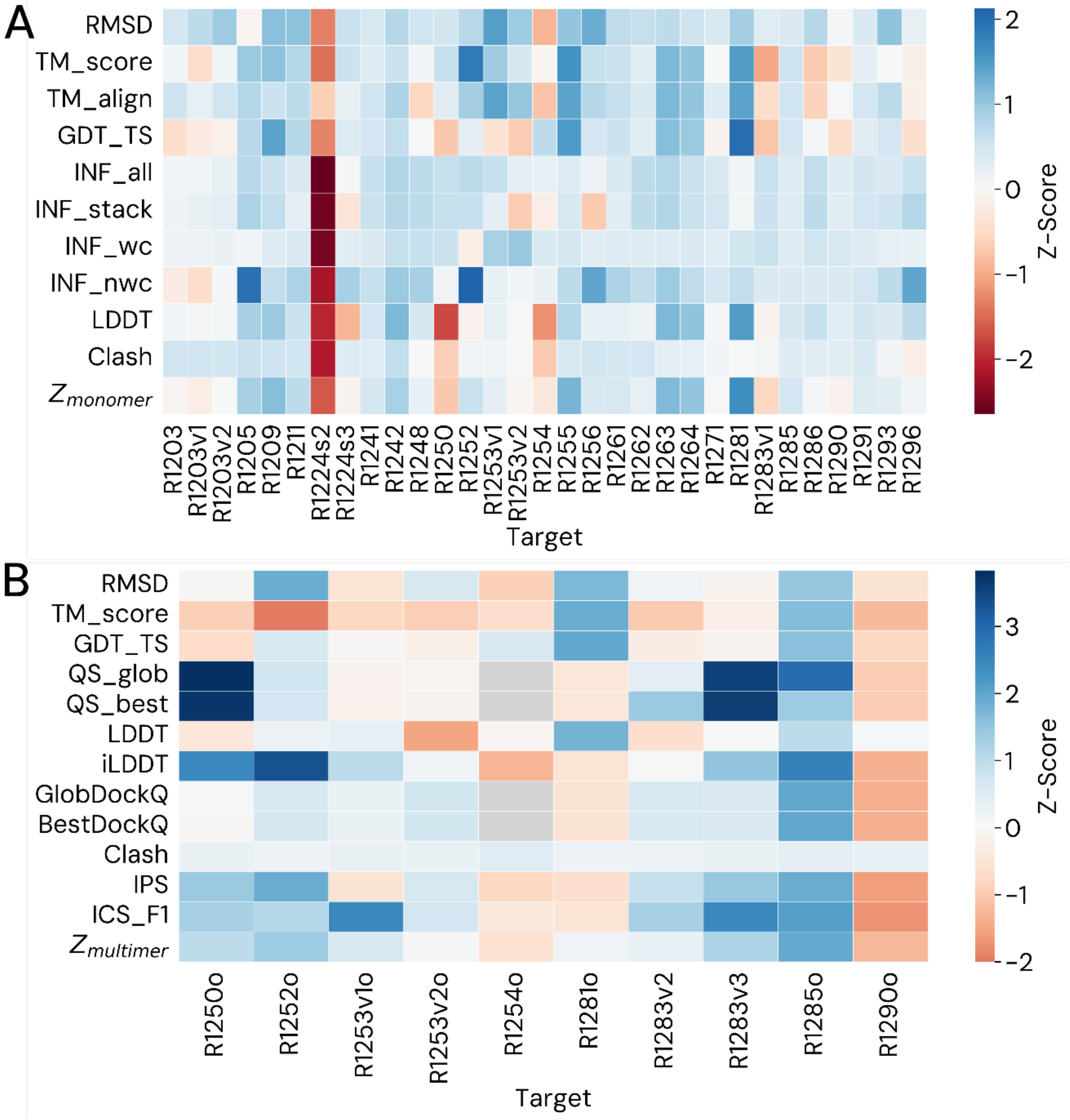


**Supplementary Figure S2. Weighted heatmap of relative model performance for the LCBio (G189) group across CASP16 targets.** (A) Monomeric RNA targets. (B) RNA–RNA multimer targets. For each target and evaluation metric, performance is expressed as a Z-score relative to the distribution across all submitted predictors, using the community mean and standard deviation reported in the official CASP16 assessments. Z-scores were sign-adjusted for metrics where lower values indicate better performance (RMSD, clash score). Individual rows correspond to structural and interface quality metrics, while columns represent targets. Composite scores ($Z_{monomer}$ and $Z_{multimer}$) summarize overall performance as weighted combinations of the underlying metrics, comprising backbone and fold accuracy for monomers and additionally interface quality for multimers. The heatmaps are displayed using a diverging color scale centred at zero, with blue indicating above-average performance and red indicating below-average performance relative to the CASP16 community. Grey cells indicate metrics where assessment data were unavailable or not computed for the target.

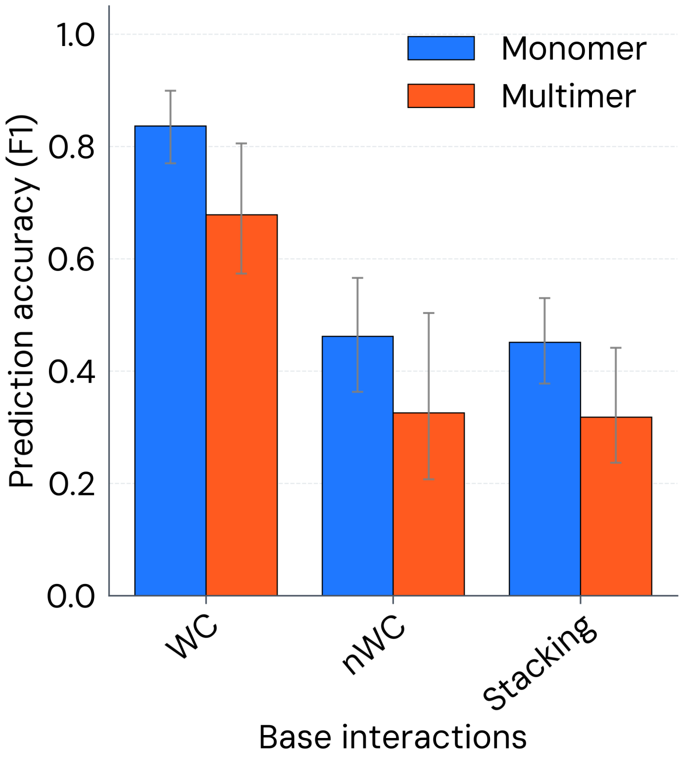


**Supplementary Figure S3. Base interaction accuracy.** Prediction accuracy (F1-score) for base interaction types: Watson–Crick base pairs, non-Watson–Crick base pairs, and base stacking interactions. Watson–Crick pairs are evaluated using exact base-pair matching. Non-Watson–Crick pairs incorporate identity, orientation, and edge classification. Stacking interactions are evaluated based on base identity and geometric agreement. Results are shown separately for monomeric and multimeric targets. Error bars represent 95% bootstrap confidence intervals accounting for non-independence among related multimer targets (see Supplementary Methods S3.3).
